# The yeast mating-type switching endonuclease HO is a domesticated member of an unorthodox homing genetic element family

**DOI:** 10.1101/2020.01.20.913210

**Authors:** Aisling Y. Coughlan, Lisa Lombardi, Stephanie Braun-Galleani, Alexandre A. R. Martos, Virginie Galeote, Frédéric Bigey, Sylvie Dequin, Kevin P. Byrne, Kenneth H. Wolfe

## Abstract

The mating-type switching endonuclease HO plays a central role in the natural life cycle of *Saccharomyces cerevisiae*, but its evolutionary origin is unknown. *HO* is a recent addition to yeast genomes, present in only a few genera. It resembles a degenerated intein fused to a zinc finger domain. Here we show that *HO* is structurally and phylogenetically related to a family of unorthodox homing genetic elements found in *Torulaspora* and *Lachancea* yeasts. These *WHO* elements integrate into the aldolase gene *FBA1*, replacing its 3’ end each time. Their structural organization is different from all known classes of homing elements. We show that a WHO protein cleaves *Torulaspora delbrueckii FBA1* efficiently and in an allele-specific manner, leading to DNA repair by gene conversion or NHEJ. The DNA rearrangement steps during *WHO* element homing are very similar to those during mating-type switching, and indicate that *HO* is a domesticated *WHO*-like element.

## Introduction

Mating-type switching is an important process in the natural life cycle of many budding yeast species. If an uninhabitable environment improves and becomes habitable, any yeast spores that germinate earlier than others will have a competitive advantage, provided that they have a backup mechanism to prevent death if they germinate too early. Mating-type switching provides this backup (Hanson and Wolfe, 2017). Yeast spores are survival capsules, and mating-type switching enables the microcolony formed from a newly-germinated spore to sporulate again after just a few cell divisions if necessary, preventing its extinction (Herskowitz, 1988). Across the phylogenetic tree of budding yeasts, mating-type switching has arisen independently at least 11 times, indicating strong natural selection in favor of switching (Krassowski et al., 2019).

In *Saccharomyces cerevisiae*, *HO* is the central gene in the mating-type switching process. It was one of the first yeast genes ever discovered, because haploid strains with a functional *HO* gene can switch their mating type and hence auto-diploidize and form visible spores, whereas *ho* mutants cannot (Winge and Roberts, 1949; Oshima, 1993). *HO* codes for an endonuclease that makes a double-strand DNA break at the mating type (*MAT*) locus, and is essential for efficient switching (Kostriken et al., 1983; Russell et al., 1986). Much of our knowledge about how eukaryotes repair double-strand DNA breaks in their chromosomes comes from studies that used HO as a model system (Haber, 2016). But despite the comprehensive functional and genetic characterization of *HO*, its evolutionary origin remains mysterious (Keeling and Roger, 1995; Haber and Wolfe, 2005; Koufopanou and Burt, 2005; Muller et al., 2007). The *HO* gene is a relatively recent evolutionary addition into the yeast genome, because it is found only in a few genera closely related to *Saccharomyces* (Butler et al., 2004; Hanson and Wolfe, 2017). The ‘three-locus’ system for mating-type switching, involving an active *MAT* locus and silent *HML* and *HMR* loci, pre-dates the origin of *HO*. An outgroup species, *Kluyveromyces lactis*, also has a three-locus system but has no *HO* gene, and employs alternative mechanisms to create a double-strand break at the *MAT* locus to initiate switching (Barsoum et al., 2010; Rajaei et al., 2014). More distantly related budding yeasts switch mating types using ‘two-locus’ flip/flop inversion systems, and again do not have an *HO* gene (Hanson and Wolfe, 2017; Krassowski et al., 2019).

As well as being a recent evolutionary innovation, HO endonuclease also has an unusual protein domain structure that begs the question of where it came from. It resembles inteins, but is not an intein itself. Inteins are mobile genetic elements that are completely protein-coding and occur as in-frame fusions within a host gene (Novikova et al., 2014). After the host gene is transcribed and translated, the intein is excised post-translationally and the host protein is assembled by protein splicing, making a peptide bond between its N- and C-terminal parts (exteins). HO has highest sequence similarity to the VDE intein of budding yeasts, which is the only intein in *S. cerevisiae* (Koufopanou and Burt, 2005; Green et al., 2018). The host gene for *VDE* is *VMA1*, which codes for a subunit of vacuolar H^+^-ATPase (Gimble and Thorner, 1992; Anraku et al., 2005). The excised VDE intein has endonuclease activity and can cleave empty (inteinless) alleles of *VMA1*, enabling the intein to spread through the population by homing – a selfish, super-Mendelian mode of inheritance (Burt and Koufopanou, 2004; Burt and Trivers, 2008). The *VMA1* genes of several species in the budding yeast family Saccharomycetaceae are polymorphic for the presence/absence of *VDE*, due to active homing and interspecies spread of the intein (Koufopanou et al., 2002; Okuda et al., 2003). Homing of *VDE* into empty alleles of *VMA1* occurs during meiosis in diploids that are heterozygotes for intein-containing and empty alleles of *VMA1* (Gimble and Thorner, 1992). The VDE protein has two domains (Moure et al., 2002): a protein splicing domain that enables the host protein Vma1 to be made, and a homing endonuclease domain that enables the *VDE* DNA sequence to home into empty alleles of *VMA1*.

Most inteins are found in bacteria and archaea, not yeasts (Poulter et al., 2007; Novikova et al., 2014; Green et al., 2018). In phylogenetic and other sequence similarity analyses, the two yeast proteins HO and VDE were found to be each other’s closest relatives (Dalgaard et al., 1997; Koufopanou and Burt, 2005; Green et al., 2018). Although HO is related to inteins, and more distantly related to other homing endonucleases in the LAGLIDADG superfamily (Chevalier and Stoddard, 2001), it is an independently expressed standalone gene, whereas inteins and other homing endonucleases are self-splicing entities embedded within their host genes (Belfort et al., 2005; Belfort, 2017). HO does not undergo protein splicing and has no exteins. HO also has a unique zinc finger domain at its C-terminus that is not present in other intein-like proteins or homing endonucleases. In *S. cerevisiae* HO, amino acid residues essential for cleavage of the *MAT* locus are located both in the zinc finger and in the endonuclease domain of the intein-like region (Meiron et al., 1995; Bakhrat et al., 2004; Bakhrat et al., 2006), and the endonuclease has a stringent requirement for zinc ions (Jin et al., 1997).

A key feature differentiating HO from true homing endonucleases is that it does not propagate its own DNA sequence – in other words, it does not home. *HO* has become a normal cellular gene, and is one of the many known examples of a mobile genetic element that has been domesticated to take on a new role (Volff, 2006; Kaiser et al., 2009; Motl and Chalker, 2009; McDowell and Meyers, 2013; Chiruvella et al., 2016; Huang et al., 2016).

However, until now it has been unclear what type of mobile element HO originated from. Here, we show that HO is related to a large and diverse family of intein-zinc finger fusion proteins (WHO proteins) that occur mostly in the yeast genus *Torulaspora*. WHO proteins are encoded by a newly discovered homing genetic element, whose genomic organization is different from all other known homing elements, and whose host is the aldolase gene *FBA1*. The similarities between WHO and HO proteins, and between the DNA rearrangement steps that occur during *WHO* element homing and mating-type switching, show how the HO-catalyzed system of mating-type switching originated.

## Results

### *WHO* genes code for a family of intein-zinc finger fusion proteins similar to HO

In the genome sequence of the type strain of *Torulaspora delbrueckii* (CBS1146; Gordon et al., 2011), we identified a cluster of five genes (*TDEL0B06670* to *TDEL0B06710*) spanning 14 kb that have sequence similarity to *HO*. We renamed these genes *WHO1* to *WHO5*, for ‘weird *HO*’ (Fig. 1A). Two of them are pseudogenes, with a single frameshift in *WHO1* and more extensive damage in *WHO5*. The *WHO* gene cluster is located downstream of the *FBA1* gene encoding fructose-1,6-bisphosphate aldolase, an enzyme that functions bidirectionally in glycolysis and gluconeogenesis (Schwelberger et al., 1989). Amino acid sequence identity among the inferred WHO proteins is unusually low for a tandem gene cluster, ranging from 55% (Who2 vs. Who4) down to 24% (Who3 vs. Who4). *T. delbrueckii* also has an *HO* gene (*TDEL0A00850*) elsewhere in its genome, orthologous and syntenic with the *HO* gene of *S. cerevisiae*. The five inferred WHO proteins have only 22-25% identity to *T. delbrueckii* HO (BLASTP *E*-values in the range 1e-6 to 3e-30).

**Figure 1.**
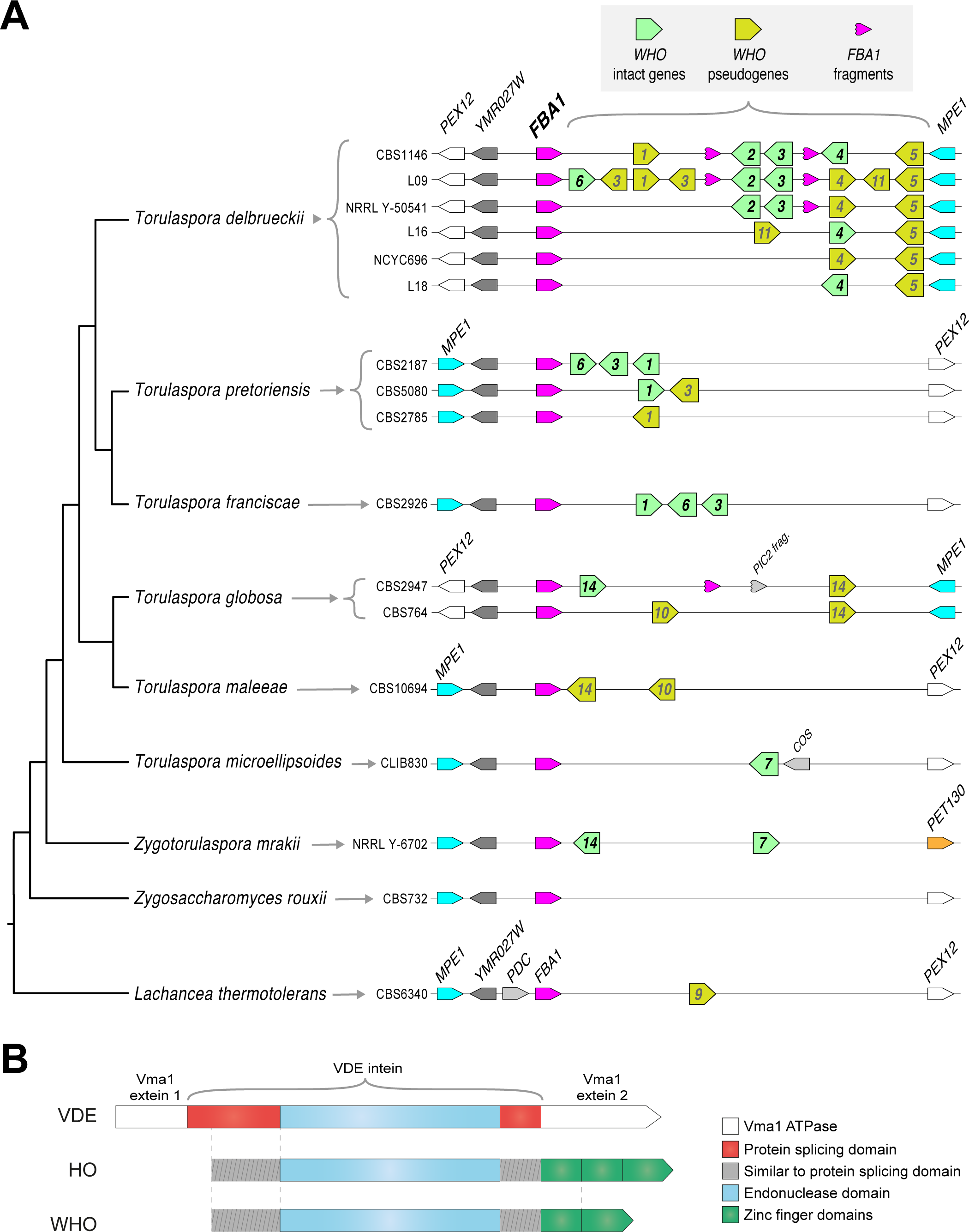
Genomic organization and domain structure of *WHO* genes. **(A)** Polymorphic clusters of *WHO* genes and pseudogenes downstream of *FBA1* in *Torulaspora* species. Multiple alleles are shown for *T. delbrueckii*, *T. pretoriensis*, and *T. globosa*. *WHO* genes are indicated by their family number. Fragments of the 3’ end of the *FBA1* gene are marked. Genomic views are schematic and not drawn to scale. The phylogenetic tree is based on Shen et al. (2018). **(B)** Domain structure of HO, VDE and WHO proteins. The protein splicing domain is formed from two regions of the protein that flank the endonuclease domain (Moure et al., 2002).

The length and content of the *WHO* gene cluster is polymorphic among natural isolates of *T. delbrueckii*. This species has a primarily haploid (haplontic) life cycle (Kurtzman, 2011), so there is only one allele per strain. We found six different allelic *WHO* cluster arrangements among 15 strains we examined, with content ranging from 2 to 9 *WHO* genes and pseudogenes (Fig. 1A; Table S1). The largest cluster (18 kb) is in strain L09, with three intact genes (*WHO6*, *WHO2*, *WHO3*) and six *WHO* pseudogenes.

As well as *WHO* genes and pseudogenes, some of the *T. delbrueckii* clusters contain one or two duplicated fragments of the 3’ end of *FBA1*, interspersed with *WHO* genes (Fig. 1A). The *FBA1* fragments are about 400 bp long (full-length *FBA1* is 1080 bp). Some *FBA1* fragments contain frameshifts or internal stop codons, and others are intact.

The WHO proteins consist of an intein-like region followed by a zinc finger domain (Fig. 1B). This structure is similar to HO, but distinct from VDE which has no zinc finger. WHO and HO are the only intein-zinc finger fusion proteins known in any organism. WHO and HO both have no exteins, and their intein domains both lack an amino acid motif that is normally found at the C-terminal end (motif G; Pietrokovski, 1994). Similar to HO, the zinc finger domains of the WHO proteins contain variable numbers (3-5) of a Cys-X-X-Cys motif. By BLAST searches, we found that the most similar zinc finger domains in other yeast proteins occur in orthologs of *S. cerevisiae* Ash1 (a regulator of *HO* transcription), which is an atypical type of GATA zinc finger domain (Scazzocchio, 2000; Munchow et al., 2002), though the level of sequence identity is low (maximally 38% over 76 residues).

### *WHO* gene/pseudogene clusters downstream of *FBA1* are common in the genera *Torulaspora* and *Lachancea*

*WHO* gene/pseudogene clusters are present downstream of *FBA1* in all five other species of the genus *Torulaspora* that we examined, and they again show within-species polymorphism in their gene content. We found 3 allelic *WHO* cluster arrangements among 9 *T. pretoriensis* strains, and 2 allelic arrangements in 2 *T. globosa* strains (Fig. 1A). A pair of *WHO* genes is also present downstream of *FBA1* in *Zygotorulaspora mrakii*, which is closely related to *Torulaspora*. There are no *WHO* genes in the next most closely related genus, *Zygosaccharomyces*.

By database searches, we found that *WHO* genes also occur in the genus *Lachancea*, but they are absent from almost every other sequenced budding yeast genome. Eleven of the 12 sequenced *Lachancea* species have clusters of *WHO* genes or pseudogenes downstream of *FBA1*, and five of these species have intact *WHO* genes (Fig. S1A). The *WHO* clusters in four *Lachancea* species also contain duplicated fragments of the 3’ end of *FBA1*. These structures in *Lachancea* are remarkably similar to the ones seen in *Torulaspora*, especially considering that these two genera are not closely related to each other, and there are no *WHO* genes in most other yeast genera. In addition, a few *Lachancea* genomes contain other *WHO* genes or pseudogenes at loci separate from *FBA1* (Fig. S1B).

In summary, *WHO* genes resemble homing endonuclease genes (Gimble, 2000). Like HO, but unlike any other homing endonucleases, the endonuclease domain is fused to a zinc finger domain. The structure of *WHO* genes, and their occurrence in clusters that are mixtures of intact genes and pseudogenes, suggests that they are part of a mobile genetic element.

### A WHO protein cleaves the *T. delbrueckii FBA1* gene in an allele-specific manner

We hypothesized that WHO proteins are homing endonucleases whose target is the *FBA1* gene. This hypothesis was motivated by two observations. First, *WHO* genes with very diverse sequences occur in tandem clusters downstream of *FBA1*, in both *Torulaspora* and *Lachancea*, suggestive of repeated integrations of different members of a mobile element family into the same target locus. Second, fragments of *FBA1* are present within the *WHO* gene clusters in both *Torulaspora* and *Lachancea* (Fig. 1A; Fig. S1A). All the *FBA1* fragments consist of only the 3’ end of the gene, and many of them begin at approximately the same position (base 670-680 in the gene sequence), which we hypothesized could indicate a possible endonuclease cleavage site in *FBA1*.

To test the hypothesis that WHO proteins target the *FBA1* gene, we carried out experiments in *S. cerevisiae* because few tools exist for genetic manipulation of *Torulaspora* or *Lachancea.* We chose *WHO6* for these experiments because it is intact, present in only a minority of the *T. delbrueckii* isolates we examined (3 of 15), and it is located at the end of the cluster closest to *FBA1* (Fig. 1A). Together, these features suggested that *WHO6* could be the most recently-inserted *WHO* gene in the array at the *T. delbrueckii FBA1* locus.

*FBA1* is an essential gene in most growth conditions (Lobo, 1984; Schwelberger et al., 1989; Boles and Zimmermann, 1993), so we reasoned that if it is the natural target of WHO endonuclease cleavage, then strains of *T. delbrueckii* that contain a *WHO6* gene should contain alleles of *FBA1* that are resistant to cleavage by Who6 endonuclease, whereas other *T. delbrueckii* strains might contain alleles that are sensitive to Who6. We constructed haploid strains of *S. cerevisiae* that contain the open reading frame (ORF) of *T. delbrueckii FBA1* (*TdFBA1*) integrated into the *ADE2* gene on chromosome XV. These strains also have the native *S. cerevisiae FBA1* gene on chromosome XI. We used two different alleles of *TdFBA1*: one from a *T. delbrueckii* isolate (strain L09) that has a *WHO6* gene downstream of it, and one from an isolate (CBS1146) that has no *WHO6* gene (Fig. 1A). We then introduced a high copy-number panARS plasmid (pWHO6-HA) on which *WHO6* was expressed from the constitutive *T. delbrueckii TDH3* promoter (Fig. 2A), with a 3xHA epitope tag at its 3’ end. As described below, we found that this plasmid induces cleavage of the allele from strain CBS1146, but not of the allele from strain L09. Hence we designated the CBS1146 allele *TdFBA1-S* (for sensitivity to cleavage by Who6), and the L09 allele *TdFBA1-R* (for resistance).

**Figure 2.**
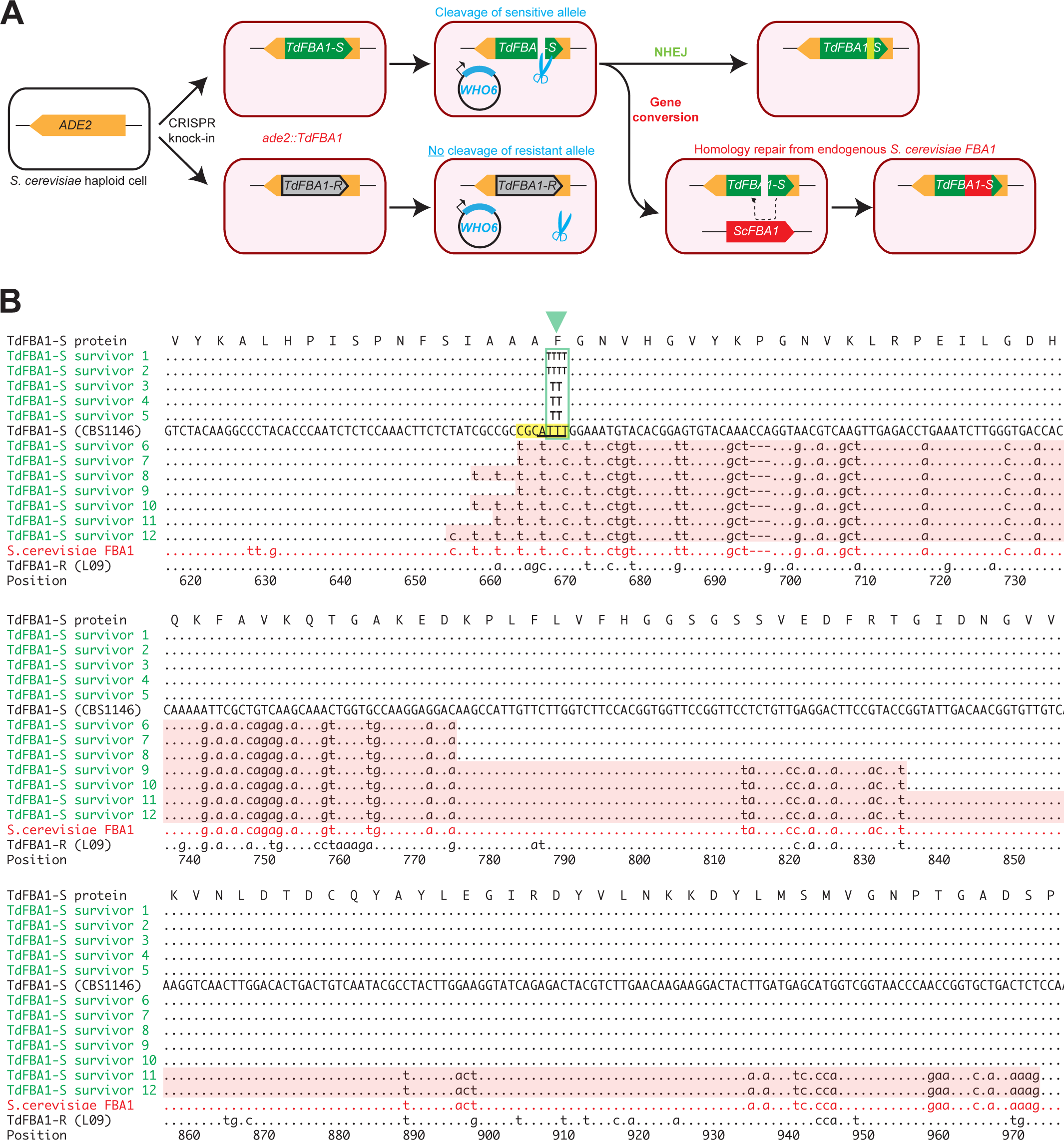
*WHO6* induces allele-specific DNA cleavage of the *T. delbrueckii FBA1* gene, with DNA repair by gene conversion or NHEJ. **(A)** Summary of the experiment. Haploid *S. cerevisiae* strains, containing a non-expressed *T. delbrueckii FBA1* ORF (*TdFBA1-S* or *TdFBA1-R* alleles) integrated at *ADE2*, were transformed multiple independent times with plasmids expressing WHO6 or WHO6-HA. In all transformations of strains containing the *TdFBA1-S* allele, the only colonies that survived expression of *WHO6* were ones in which *TdFBA1-S* underwent DNA cleavage and repair by gene conversion or imprecise NHEJ, changing its sequence and making it resistant to further cleavage. **(B)** Gene conversion and imprecise NHEJ events in *TdFBA1-S*. The reference sequence (uppercase) shows the 3’ end of the *TdFBA1-S* allele from *T. delbrueckii* strain CBS1146. Survivors 1-5 are transformants in which *TdFBA1-S* was cleaved by Who6 and repaired by imprecise NHEJ near position 668 (green box and triangle; survivors 1 and 2 have a 1-bp insertion, and survivors 3-5 have a 1-bp deletion, relative to the sequence TTT in the reference). Survivors 6-12 are transformants in which *TdFBA1-S* was cleaved by Who6 and partially overwritten by gene conversion with the endogenous *S. cerevisiae FBA1* gene. Gene conversion regions are highlighted with pink backgrounds. The *TdFBA1-R* allele from *T. delbrueckii* strain L09, which is the natural host of *WHO6*, is also shown; this allele acquired no sequence changes among 10 independent pWHO6 transformants examined. A putative Who6 recognition site (yellow) and cleavage site with 4-bp 3’ overhang (underlined) are marked. Survivors 10 and 11 are from transformations with pWHO6-HA; all other survivors are from transformations with pWHO6. The complete *TdFBA1* gene was sequenced from all transformants but only positions 616 to 975 are shown; there were no changes outside this region.

Our experiment is similar to one carried out by Moore and Haber (1996) who overexpressed HO so that it continually cleaved the *MAT* locus in haploid *S. cerevisiae* cells that had no *HML/HMR* loci. They found that the only cells that survived HO overexpression were ones in which inaccurate DNA repair ligated the chromosome back together but modified the target site sequence in such a way that HO could no longer cleave it, because chromosomes with accurate repairs were re-cleaved by HO. Similarly, in our experiment, the only haploid *cerevisiae* cells containing *TdFBA1-S* that survived overexpression of Who6 were ones in which the *TdFBA1-S* sequence became modified, either by gene conversion or by imprecise non-homologous end joining (NHEJ) (Fig. 2A).

By genome sequencing, we found that in two independent experiments where pWHO6-HA was introduced into *S. cerevisiae* strains containing *TdFBA1-S*, this *T. delbrueckii FBA1* allele was modified by gene conversion with the native *S. cerevisiae FBA1* gene. In contrast, no gene conversion was seen when pWHO6-HA was introduced into *S. cerevisiae* strains containing the *TdFBA1-R* allele (two independent transformants), nor in control transformations in which a similar plasmid expressing 3xHA-tagged Green Fluorescent Protein (pGFP-HA) was transformed into *S. cerevisiae* strains containing *TdFBA1-S* or *TdFBA1-R.* Apart from the gene conversions at the *TdFBA1-S* transgene, no other nucleotide changes were detected in the genomes of the strains transformed with pWHO6-HA, and the native *S. cerevisiae FBA1* gene remained unchanged in this experiment.

To verify that the gene conversions were not caused by the 3xHA tag, we carried out additional experiments in which *S. cerevisiae* strains containing either *TdFBA1-S* or *TdFBA1-R* were each independently transformed 10 times with either pWHO6 (a plasmid expressing Who6 with no 3xHA tag) or pGFP-HA as a control. A single colony was chosen from each transformation, and its *ade2::TdFBA1* locus was amplified by PCR and sequenced. Dramatically fewer colonies were obtained from the transformations of pWHO6 into the *TdFBA1-S* strain than in any of the other three combinations of plasmid (pWHO6 or pGFP-HA) and allele (*TdFBA1-S* or *TdFBA1-R*). Evidence of cleavage of the *T. delbrueckii FBA1* ORF was observed in all 10 independent transformants of pWHO6 into the *TdFBA1-S* strain: 5 transformants showed gene conversion between *TdFBA1-S* and the native *S. cerevisiae FBA1* gene, and the other 5 had single-nucleotide insertions or deletions in *TdFBA1-S* consistent with cleavage and repair by imprecise NHEJ (Fig. 2B). In contrast, no sequence changes at *ade2::TdFBA1* were detected in the 10 independent transformants of pWHO6 into the *TdFBA1-R* strain, nor (as expected) in any of the 20 transformants with pGFP-HA.

From these experiments we conclude that the *T. delbrueckii* isolate (L09) that contains the *WHO6* gene also contains, 588 bp upstream, an allele of *FBA1* (*TdFBA1-R*) that is resistant to cleavage by the Who6 endonuclease. It can therefore stably maintain this endonuclease gene in its genome. In contrast, an isolate (CBS1146) that has no *WHO6* gene contains an *FBA1* allele (*TdFBA1-S*) that is sensitive to cleavage by Who6.

### The WHO endonuclease cleavage site in *FBA1*

Similar to the HO overexpression survivors in Moore and Haber (1996), our transformants that contained the sensitive *TdFBA1-S* allele of *T. delbrueckii FBA1* and survived overexpression of Who6 have acquired mutations that can be inferred to have damaged the Who6 recognition or cleavage sites, making the cells resistant to Who6. These mutations enable us to identify the approximate location of the Who6 recognition and cleavage sites in *S. delbrueckii FBA1*.

The five transformants showing evidence of imprecise NHEJ (survivors 1-5 in Fig. 2B) each sustained a single 1-bp insertion or deletion in the *TdFBA1-S* sequence, at the same site in the gene (positions 667-669: TTT→TTTT, or TTT→TT). Like other LAGLIDADG endonucleases, HO and VDE both make a staggered double-strand break with 4-bp 3’ overhangs when they cleave DNA, and they have large (∼24 bp) degenerate recognition sequences that span the cleavage site (Nickoloff et al., 1990; Gimble and Thorner, 1993; Taylor et al., 2012). We therefore infer that the overhang made by Who6 must include some or all of the TTT sequence centered on position 668 (Fig. 2B). The HO recognition site at the *MAT* locus is moderately conserved between *S. cerevisiae* and other budding yeasts, and has at its core the sequence CGC<u>AACA</u>, where the 4-bp overhang is underlined. By analogy, we suggest that the core of the Who6 recognition site in the sensitive *TdFBA1-S* allele of *T. delbrueckii FBA1* is CGC<u>ATTT</u> (positions 663-669). In the resistant *TdFBA1-R* allele, 3 of these 7 bases are different (Cag<u>cTTT</u>). In *S. cerevisiae FBA1*, which is also resistant to cleavage by Who6, 3 of 7 bases are different (tGC<u>tTTc)</u> (Fig. 2B).

The seven transformants showing evidence of gene conversion (survivors 6-12, including 2 from pWHO6-HA transformations and 5 from pWHO6 transformations) each replaced a section of the *T. delbrueckii FBA1-S* sequence with the corresponding section of the *S. cerevisiae FBA1* gene from chromosome XI (Fig. 2B). The *T. delbrueckii* and *S. cerevisiae* genes have 84% nucleotide sequence identity overall. The gene conversion tracts are asymmetrical: they extend rightwards (towards the stop codon of *FBA1*) from the cleavage site for 106–306 bp, whereas they extend leftwards for only 5–14 bp. This asymmetry suggests that one side of the WHO-induced double-strand break is more active in recombination than the other. Similarly, during mating-type switching in *S. cerevisiae*, the DNA on one side of the HO-induced break (the Z-side) participates in exchange with *HML/HMR*, whereas DNA on the Y-side remains inert until it is eventually clipped off (Lee and Haber, 2015).

In summary, the pattern of NHEJ events and gene conversions in *S. cerevisiae* cells that contain *TdFBA1-S* and survive continuous expression of a WHO protein is very similar to the pattern in cells that contain *MAT* and survive continuous expression of HO (Moore and Haber, 1996). We infer that WHO endonucleases cleave *FBA1* genes at approximately base 668, which is slightly upstream of the 5’ ends of many of the *FBA1* fragments seen in *Torulaspora* and *Lachancea* species. The core of the putative recognition site of a WHO endonuclease is also similar to that of HO, with the sequence 5’-CGC-3’ adjacent to the overhang.

### Repeated homing continually replaces the 3’ end of *FBA1* and builds *WHO* clusters

The *TdFBA1-S* and *TdFBA1-R* alleles have only 85% nucleotide sequence identity downstream of base 668, which is remarkably low for two alleles from the same species. In contrast, they have 99% identity upstream of this position. More generally, among the full-length *FBA1* genes of the 15 *T. delbrueckii* isolates we analyzed, nucleotide sequence diversity is much higher in the 3’ part of the gene (Fig. 3A). Moreover, phylogenetic trees constructed from the 5’ and 3’ parts of *TdFBA1* (upstream and downstream of position 668) have contradictory topologies (Fig. 3B). The heterogeneous evolution of the two ends of *FBA1* in *T. delbrueckii*, and the presence of 3’ *FBA1* fragments in its genome, suggest a mechanism for how the homing genetic element containing *WHO* genes operates.

**Figure 3.**
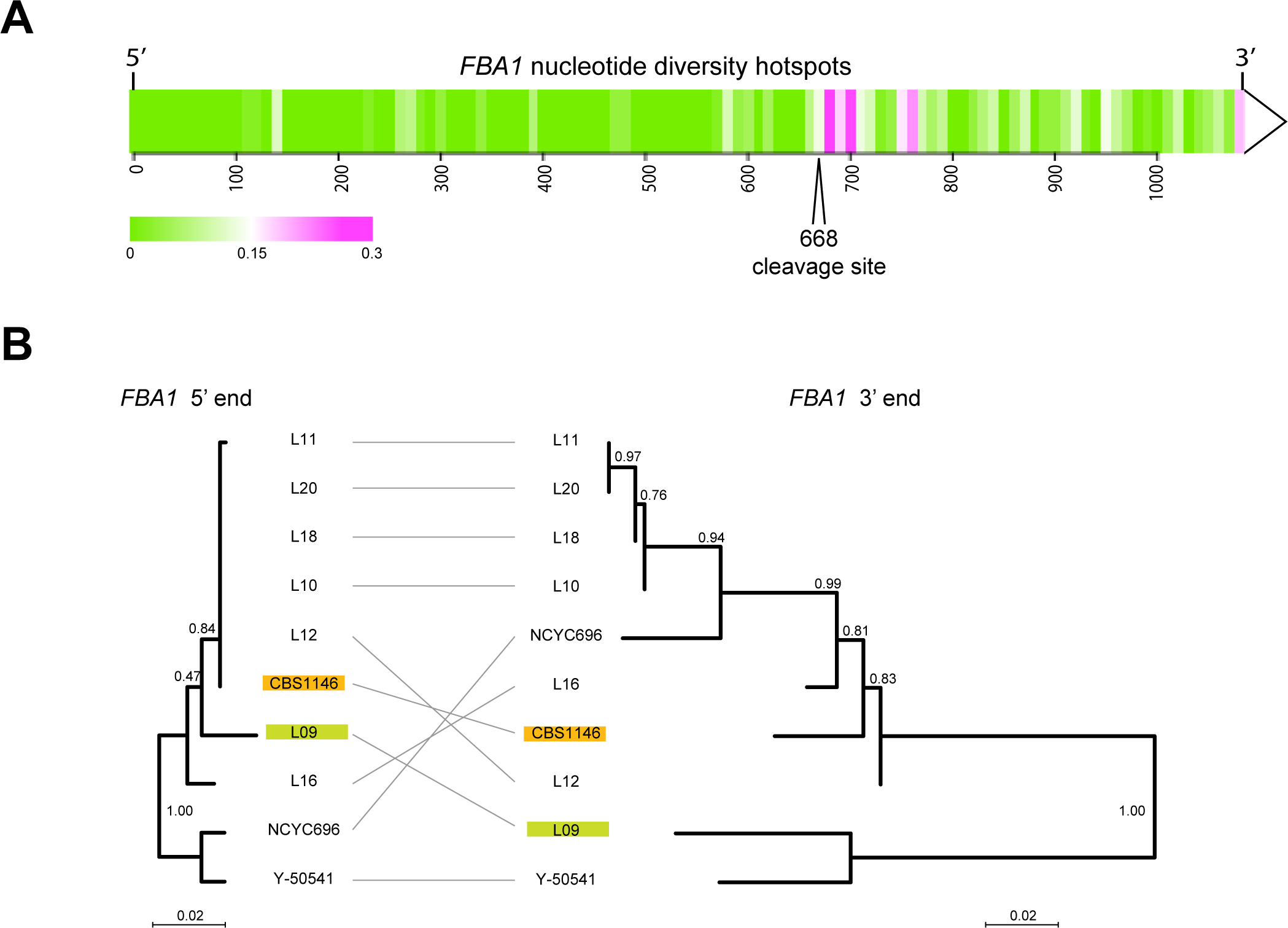
Different evolutionary dynamics of the 5’ and 3’ parts of *T. delbrueckii FBA1.* **(A)** Heatmap showing regions of nucleotide sequence diversity (*π*) among 15 sequenced *FBA1* alleles from *T. delbrueckii* isolates, plotted in 10-bp windows. **(B)** Inconsistency of phylogenetic trees obtained from the 5’ and 3’ ends of *T. delbrueckii FBA1* alleles (bases 1-667, and 669-1083, respectively). The positions of the alleles from strains CBS1146 (*TdFBA1-S*) and L09 (*TdFBA1-R*) are highlighted. The trees are drawn to the same scale and were generated using PhyML as implemented in Seaview v4.5.0 (Gouy et al., 2010) with default parameters. Bootstrap support from 1000 replicates is shown.

We propose that *WHO* elements home into the *FBA1* locus by using a mechanism that involves replacing the 3’ end of *FBA1*, thereby converting a sensitive *FBA1* allele into one that is resistant to the particular WHO protein encoded by the element (Fig. 4A, left column). During meiosis in a heterozygous diploid cell, the WHO protein encoded by the donor allele cleaves the sensitive full-length *FBA1* gene of the recipient allele at position 668. The double-strand break is then repaired by using the *WHO*-containing chromosome as a template. The 3’ region of the cleaved full-length *FBA1* gene interacts with the *FBA1* fragment in the template, resulting in incorporation of the *WHO* gene into the previously empty allele. After homing, the recipient chromosome contains a *WHO* gene located between a resistant full-length *FBA1* gene (a chimera of the recipient’s previous 5’ end and a copy of the donor’s 3’ end) and a new *FBA1* fragment formed from the recipient’s previous 3’ end. In this model, the *FBA1* fragment downstream of the donor’s *WHO* gene is an essential part of the *WHO* element because it provides a region of homology that acts as a recombination site (Fig. 4A). The fragment is not part of the expressed *FBA1* gene so it does not need to maintain an open reading frame.

**Figure 4.**
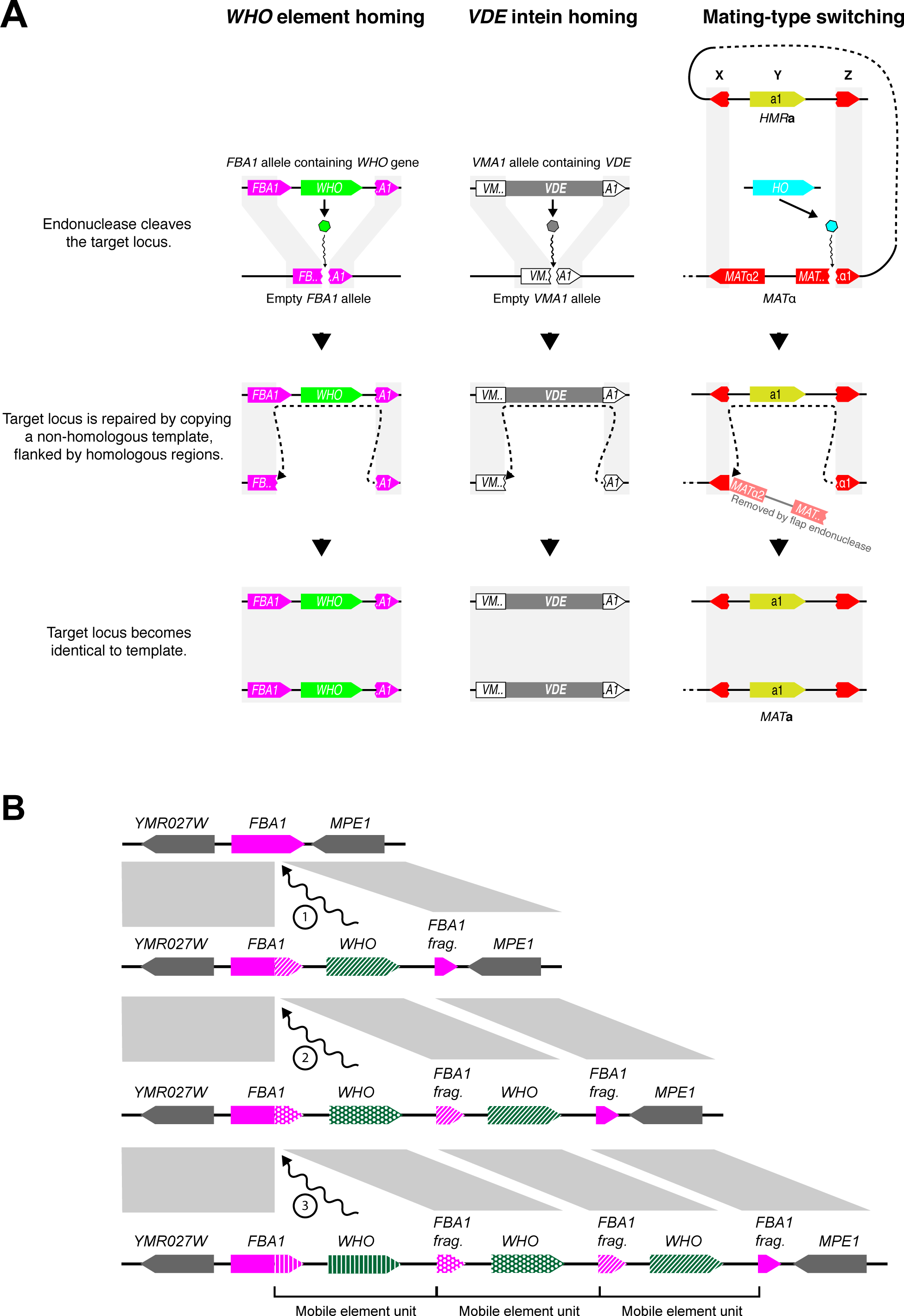
(A) Similarity of mechanisms of action of *WHO*, *VDE* and *HO*. The mechanism we propose for *WHO* elements integrating at *FBA1* is compared to the known mechanisms for *VDE* integrating into *VMA1* and for *HO*-mediated switching of the *MAT* locus (Gimble and Thorner, 1992; Lee and Haber, 2015). *WHO* and *VDE* homing occur between allelic chromosomes in a diploid cell, whereas mating-type switching occurs between *MAT* and *HML/HMR* loci in a haploid cell. Gray rectangles indicate regions of sequence identity. The column on the right shows mating-type switching from *MAT*α to *MAT***a** in *S. cerevisiae*. Switching from *MAT***a** to *MAT*α occurs by an identical mechanism; the core of the HO recognition site (CGCAACA) is the first 7 nucleotides of the Z region, which is present in both of the *MAT* alleles even though it is part of the *MAT*α1 gene sequence. The *HO* gene is on a different chromosome than *MAT*-*HML-HMR*. **(B)** Model for *WHO* cluster formation by successive integration of *WHO* elements. Every time a *WHO* element integrates into the locus, the 3’ end of the full-length *FBA1* gene is replaced. The previous 3’ end is pushed rightwards, together with any older *WHO* genes, after which they can decay into pseudogenes. The complete *WHO* mobile element unit consists of a *WHO* gene and the upstream 3’ end of *FBA1* which confers resistance to it.

While the modified *FBA1* is now resistant to the newly acquired *WHO* gene, it may still be sensitive to other *WHO* genes. Repeated homing of multiple different *WHO* elements into the same chromosome will build tandem clusters of *WHO* genes and *FBA1* fragments, with the most recent elements being located closest to the full-length *FBA1* gene (Fig. 4B). Each homing event replaces the 3’ end of the full-length *FBA1* gene with the sequence from a different allele, causing its rapid evolution and discordant phylogenies. Over time, the *WHO* genes and *FBA1* fragments can decay into pseudogenes or become deleted, because they are not required for the aldolase function of *FBA1.* The functional unit of a *WHO* element can be defined as a *WHO* gene and the resistance-conferring *FBA1* 3’ region upstream of it (Fig. 4B).

In summary, *T. delbrueckii WHO* genes are part of a homing genetic element that targets *FBA1.* Our model for its mechanism of action explains how *WHO* clusters and *FBA1* fragments are formed, and the unusual chimeric mode of evolution of *FBA1*.

### Phylogeny of *WHO* genes shows an *FBA1*-associated backbone and multiple transpositions to other genomic loci

We investigated the phylogenetic relationship among *WHO*, *HO*, and *VDE* genes, using amino acid sequences inferred from intact genes and from some of the less-damaged *WHO* pseudogenes. In view of the high divergence among the sequences, the tree topology may not be fully accurate, but it permits identification of approximately 14 families of *WHO* genes (Fig. 5). The *WHO* families form a monophyletic group, separate from *HO* and *VDE*. Most of the families are either *Torulaspora*-specific or *Lachancea-*specific, indicative of recent gene duplications within each genus. The *WHO2*, *WHO4*, *WHO5* and *WHO11* families are specific to *T. delbrueckii* so they must be young. Overall, the tree indicates a dynamic history of extensive *WHO* gene duplication and frequent formation of pseudogenes, consistent with the ‘cycle of degeneration’ expected for a homing genetic element (Burt and Koufopanou, 2004).

**Figure 5.**
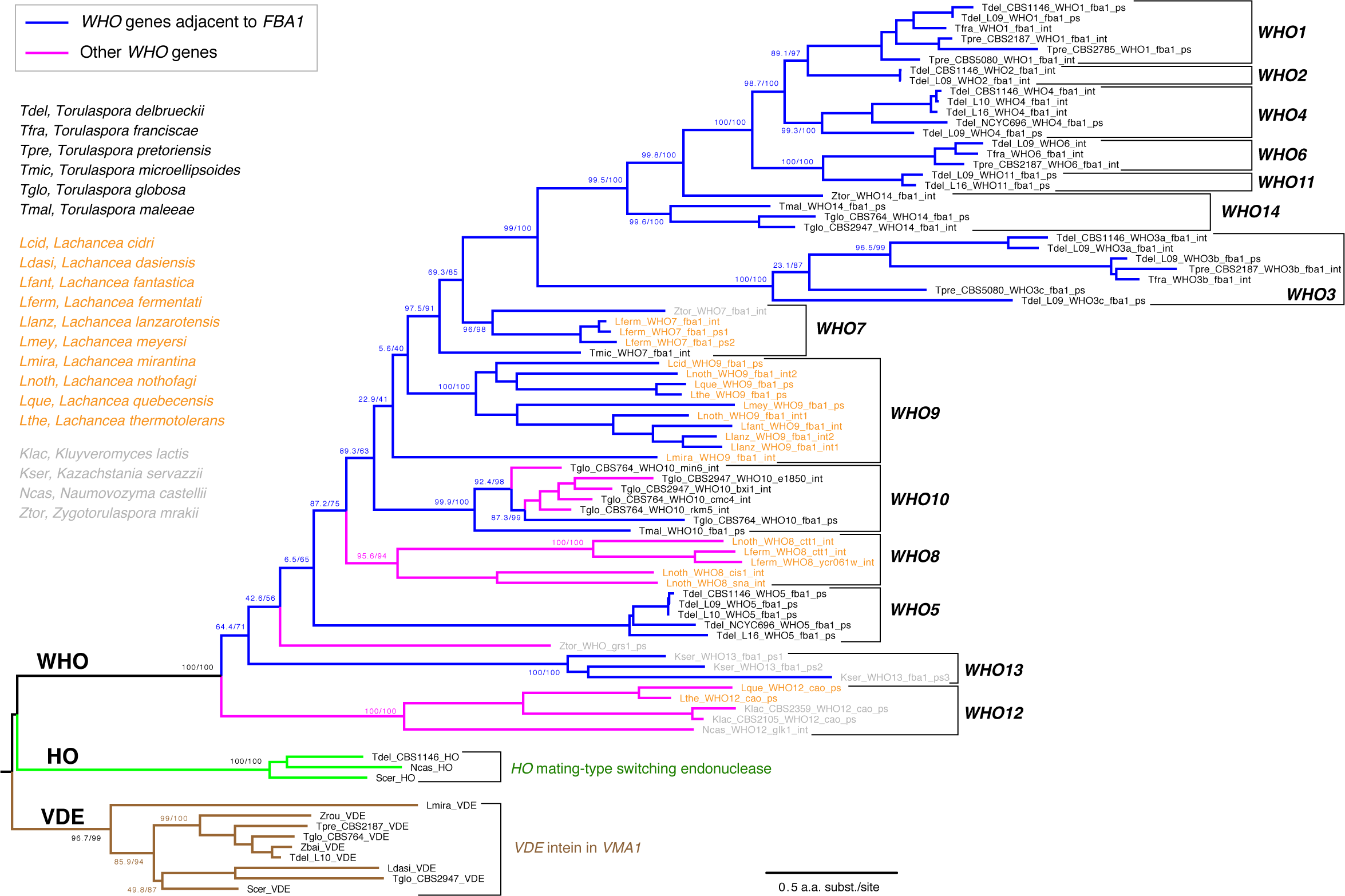
Families of *WHO* genes and their phylogenetic relationship to *HO* and *VDE.* Blue branches indicate *WHO* genes that are located at the *FBA1* locus, and magenta branches indicate *WHO* genes that are not beside *FBA1*. 14 *WHO* families are marked by brackets. Individual *WHO* gene names are colored by their source genus (black, *Torulaspora*; orange, *Lachancea*; gray, other genera). *WHO* gene names indicate the source species and strain number (if multiple strains were analyzed), *WHO* family (in uppercase), the name of a neighboring gene in the genome (in lowercase), and the suffix “int” for intact *WHO* genes or “ps” for *WHO* pseudogenes. Protein sequences were aligned using MUSCLE and filtered with Gblocks as implemented in Seaview v4.5.0 (Gouy et al., 2010). Badly degraded pseudogenes (relics) were not included. The tree was constructed by maximum likelihood using IQ-TREE v1.6.12 (Trifinopoulos et al., 2016), utilizing the built-in model finder option. Numbers on branches show support values from SH-aLRT and 1000 ultrafast bootstraps, separated by a slash (Trifinopoulos et al., 2016). The tree was rooted using VDE because WHO and HO share a zinc finger domain.

Although most *WHO* genes and pseudogenes are located downstream of *FBA1* genes (blue branches in Fig. 5), a few of them are not. It is striking that these non-*FBA1*-associated genes fall into a small number of clades (magenta branches in Fig. 5). The *WHO* genes in these clades seem to have lost their target specificity for *FBA1* and transposed to other places in the genome, and several of these *WHO* genes are intact. Most notably, the *WHO10* family includes five intact genes from *T. globosa* that are located at five different places in the genomes of the two isolates we sequenced (Fig. 6). There is a *WHO10* pseudogene beside *FBA1* in one *T. globosa* isolate and the sister species *T. maleeae* (Fig. 1A), indicating that *WHO10* was originally associated with *FBA1*. Another *WHO* clade unlinked to *FBA1* is *WHO8*, which is present only in two species of *Lachancea* (Fig. 5).

**Figure 6.**
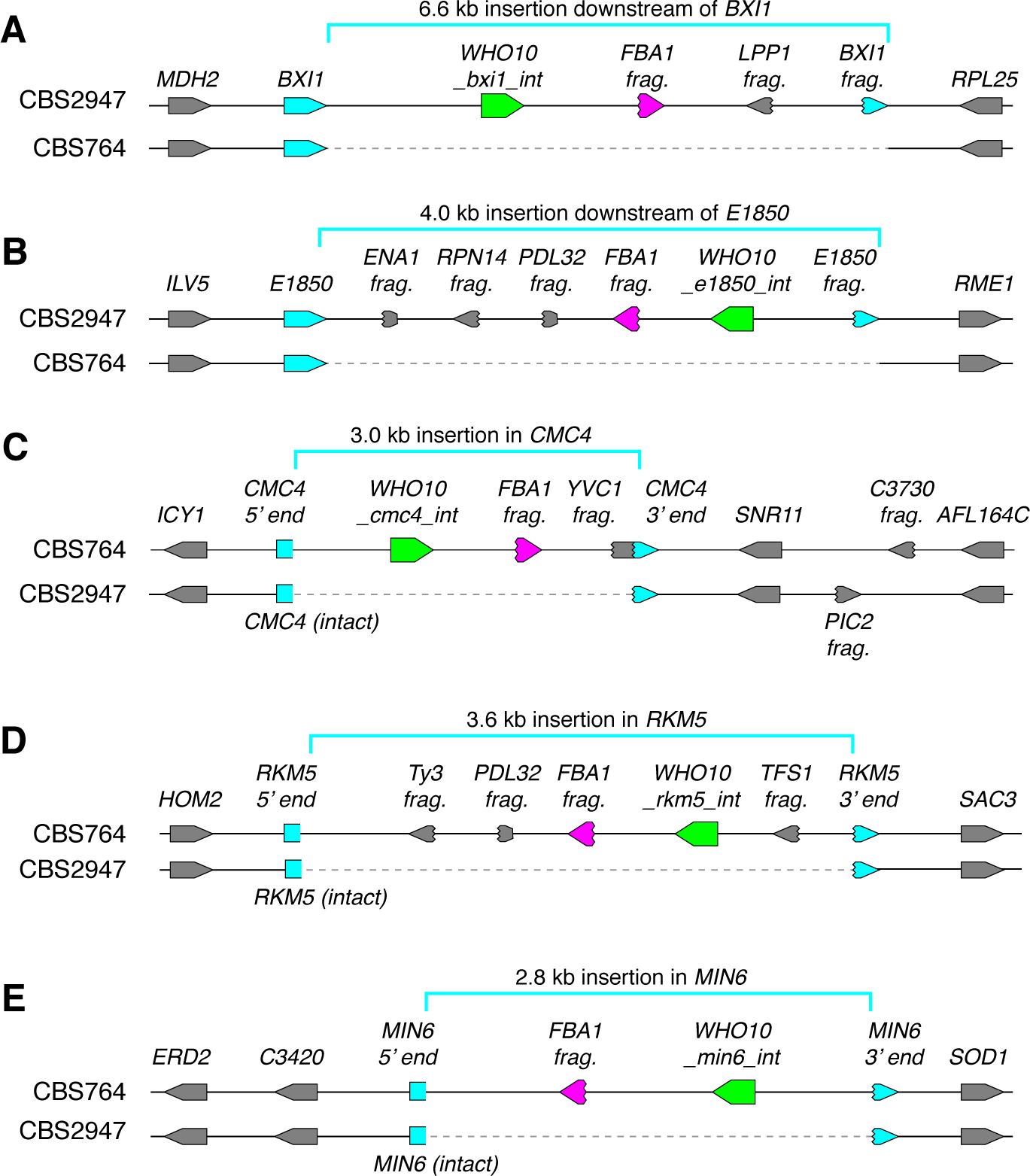
*WHO10* is an active mobile genetic element in *T. globosa* and integrates into loci other than *FBA1*. Each panel shows a pair of allelic regions from *T. globosa* strains CBS764 and CBS2947. Intact *WHO10* genes are shown in green, *FBA1* fragments in magenta, and host genes in blue. **(A-E)** Five strain-specific insertions of *WHO10* elements into different host loci. At the *BXI1* and *E1850* loci (A,B), the 3’ end of the host gene became duplicated, whereas at the *CMC4*, *RKM5* and *MIN6* loci (C-E) the host gene was disrupted by the insertion and no part of it became duplicated. All the integrations contain a fragment of *FBA1* immediately downstream of *WHO10*, and most also contain fragments of other genes. Genes are named after their *S. cerevisiae* orthologs where possible. *E1850* is the *T. globosa* ortholog of *T. delbrueckii TDEL0E01850*, a gene with no homolog in baker’s yeast.

We detected only a few *WHO* sequences in species other than *Torulaspora* (or *Zygotorulaspora*) and *Lachancea* in BLAST searches against the NCBI database, which includes genome sequences from hundreds of yeast species including members of almost every genus in the family Saccharomycetaceae (Shen et al., 2018). Thus the *WHO* family has a very limited phylogenetic distribution, and occurs mostly in two genera that are not sisters of each other (Shen et al., 2018). The few *WHO* sequences outside *Torulaspora* and *Lachancea* all lie in the *WHO13* and *WHO12* families, which are outgroups to the other *WHO* families (Fig. 5), and most of them are pseudogenes. *WHO13* is *FBA1-*associated but *WHO12* is not. The *WHO13* sequences are all pseudogenes and were detected only in a small clade of *Kazachstania* species, downstream of *FBA1*. Interestingly, these species have an intron in *FBA1* at precisely the inferred WHO endonuclease cleavage site (Fig. S2), and this is the only intron in any budding yeast *FBA1* gene. In other eukaryotes, evolutionarily novel introns are gained at sites of double-strand DNA breakage (Li et al., 2009).

The position of *WHO12* as an outgroup to all the other *WHO* families (Fig. 5) raises the possibility that the common ancestor of all the *WHO* families might have used a different gene as its original target, before changing target to *FBA1* after the *WHO12* family separated from the others. The only intact *WHO12* gene occurs in *Naumovozyma castellii*, where it is located beside *GLK1* (galactokinase). We sequenced the genomes of four strains of this species and found two with intact *WHO12*, and two with frameshifted *WHO12* pseudogenes. There was no structural polymorphism or high sequence divergence at this locus in *N. castellii*, and no *GLK1* gene fragments, so no evidence of active homing. The other *WHO12* sequences are pseudogenes in *K. lactis* (Fabre et al., 2005) and two *Lachancea* species (Fig. S1), beside a *CAO* copper amine oxidase gene in all three cases. The *K. lactis WHO12* pseudogene lies between full-length *CAO* and a damaged fragment of *CAO.* It is therefore possible that *WHO12* is an active element targeting *CAO*, and that *CAO* pre-dates *FBA1* as the target of the whole *WHO* superfamily, but given the rarity of *WHO12* it seems more likely that *FBA1* is the ancestral target and *WHO12* is a family that lost its specificity for *FBA1* more recently, similar to *WHO10*.

In summary, the phylogenetic tree indicates that *WHO* genes have been located beside *FBA1* throughout most of their diversification. The *WHO* genes that are not now located beside *FBA1* are descended from *FBA1*-linked ancestors. The patchy taxonomic distribution of *WHO* genes suggests that they are native to the genus *Torulaspora* and/or *Lachancea* and have probably been transmitted between these genera. The split between the zinc finger-containing proteins HO and WHO pre-dates the diversification of the *WHO* families.

### Recent transposition of *WHO10* genes in *Torulaspora globosa* confirms the mobility of *WHO* elements

The *WHO10* family has become amplified in *T. globosa*. In the two strains that we sequenced, strain-specific insertions of intact *WHO10* genes are present at five loci not linked to *FBA1* (Fig. 6). The five Who10 proteins have only 72–80% amino acid sequence identity to one another. The *T. globosa WHO10* genes are located within regions of inserted DNA 2.8–6.6 kb long. In two cases, the inserted DNA includes a duplicated fragment of the 3’ end of a host gene (*BXI1* and *E1850*) at one end, so that the host gene was not disrupted (Fig. 6A,B). These duplications resemble the *FBA1* fragments seen downstream of *FBA1* in many species (Fig. 1A). In the other three cases, the *WHO10-*containing insertion interrupts a host gene without forming a duplication, probably inactivating it (Fig. 6C,D,E).

All five *T. globosa* insertions include an *FBA1* fragment immediately downstream of the *WHO10* gene, even though the insertions are not near the *FBA1* locus. Some insertions also contain fragments of various other genes, mostly from their 3’ ends (Fig. 6). The structure and variable location of the DNA insertions indicate that *WHO10* genes are part of a mobile genetic element that is active in *T. globosa*. The mobile element consists of *WHO10* and the gene fragments, which may be molecular fossils of previous host sites into which the element inserted.

The genomic evidence indicates that most *WHO* genes function as part of a homing genetic element that targets the *FBA1* locus (Fig. 4). However, in *T. globosa* the *WHO10* family has lost its specificity for *FBA1* and become a more general mobile element rather than a homing element, resulting in the proliferation of intact *WHO10* genes to multiple other sites in the genome. How the *WHO10* genes become integrated into the non-*FBA1* sites in *T. globosa* is not clear, because there are no homologous flanking sequences to guide integration of *WHO10* into a double-strand break.

## Discussion

We have shown that *WHO* elements are homing genetic elements in the budding yeast genera *Torulaspora* and *Lachancea*, that primarily target the aldolase gene *FBA1*. They have diversified into a large family with very divergent endonuclease genes. Our model proposes that *WHO* elements home into sensitive alleles of *FBA1* by using a duplicated fragment of the 3’ end of *FBA1* as a second region of homology downstream of the *WHO* gene (Fig. 4A). Homing replaces the 3’ end of *FBA1*, making it resistant to cleavage by the element’s WHO endonuclease. The DNA manipulation steps in *WHO*’s homing mechanism are identical to those during *VDE* intein homing into *VMA1*, but the structures of the elements are different (Fig. 4A). Resistance to endonuclease cleavage in *FBA1* comes from allelic sequence differences, whereas resistance in *VMA1* comes directly from interruption of the cleavage site by the *VDE* element (Gimble and Thorner, 1992).

*FBA1* can be described as the host gene for the *WHO* element, even though the element lies downstream of *FBA1* rather than interrupting it. This structural organization makes *WHO* elements different from the two currently recognized classes of homing genetic elements, which are inteins and intron-encoded homing endonucleases (Belfort et al., 2005; Belfort, 2017). In both of these other classes the homing element is a self-splicing entity, transcribed as an internal part of the host gene, that must be removed to make the mature host protein. In contrast, *WHO* genes are transcribed independently of *FBA1* (some of them are in the opposite orientation to *FBA1*; Fig. 1A), and *FBA1* will remain functional even if the *WHO* gene becomes a pseudogene. *WHO* elements therefore constitute a third structural class of homing element, and the only one with a propensity to form clusters.

The reason why *WHO* elements chose *FBA1* as their host gene is probably because aldolase is absolutely required for spore formation, due to its role in gluconeogenesis (Dickinson and Williams, 1986). Meiosis and sporulation require cells to be grown on a non-fermentable carbon source such as acetate, and in these conditions gluconeogenesis is necessary to make the glucose monomers used for synthesis of the polysaccharide layers of the spore wall, a late stage in the meiosis-sporulation pathway (Neiman, 2005). *S. cerevisiae fba1* mutants cannot make spores (Lobo, 1984; Dickinson and Williams, 1986) but they should not be blocked in meiosis, which is when *WHO* element homing is expected to occur. It is unlikely that the *FBA1* genes in either the donor or recipient chromosome can be transcribed at the same time as DNA cleavage and recombination is occurring during homing. By temporarily inactivating *FBA1*, the *WHO* element may be able to delay the cell from progressing from meiosis into sporulation until homing has finished. Homing is likely to be a slow process, because mating-type switching takes more than an hour (Lee and Haber, 2015).

The DNA manipulation steps during HO-catalyzed mating-type switching strongly resemble the mechanisms of homing of *WHO* elements and the *VDE* intein (Fig. 4A). Together with the sequence similarity among the three proteins, it indicates a shared evolutionary origin of the three processes. While the mechanisms of *WHO* and *VDE* homing are similar, the mechanism of HO action at the *MAT* locus has diverged from them in two critical ways. First, the *HO* gene is not part of the template used for DNA repair. Second, switching occurs in haploids, whereas homing occurs in diploids during meiosis. There is no homologous chromosome for the cleaved *MAT* locus to interact with, so instead it interacts with *HML* or *HMR*.

Our results finally illuminate the origin of HO endonuclease. Based on the fact that WHO and HO share features that are otherwise unique – the presence of a zinc finger domain and the absence of exteins – we propose the following evolutionary model (Fig. 7). (1) An intein from a bacterial source invaded the *VMA1* gene of an early budding yeast species to become *VDE*. (2) *VDE* subsequently duplicated and mis-homed into a zinc finger protein gene, close to the 5’ end, to make a fusion gene that was the common ancestor of *WHO* and *HO*. The zinc finger directed the endonuclease to new target gene(s) in the genome. (3) The fusion gene became located between a target gene and a duplicated fragment of the target gene, forming a proto-*WHO* element. The target gene may have been *FBA1*, or possibly a different, unknown, gene. (4) The proto-*WHO* element diversified and spread through yeast populations and into additional species, with *FBA1* as its main target. Occasional mis-homing events spread the element into new targets such as the *WHO10* locations in *T. globosa.* (5) At an early stage of diversification, a WHO endonuclease developed an ability to cleave *MAT*α1, in a species that already contained a three-locus *MAT/HML/HMR* mating-type switching system, and became domesticated as *HO*. The boundary between the Y and Z regions of the *MAT* locus, which was previously variable among species, became permanently fixed at the site where the endonuclease cleaved *MAT*α1 (Fig. 4A; Hanson and Wolfe, 2017). HO’s origin from a homing element may help explain some unusual properties of this protein *in vitro*, such as its extreme catalytic inefficiency and its ability to attach to both ends of linear DNA molecules, forming loops visible by electron microscopy (Jin et al., 1997).

**Figure 7.**
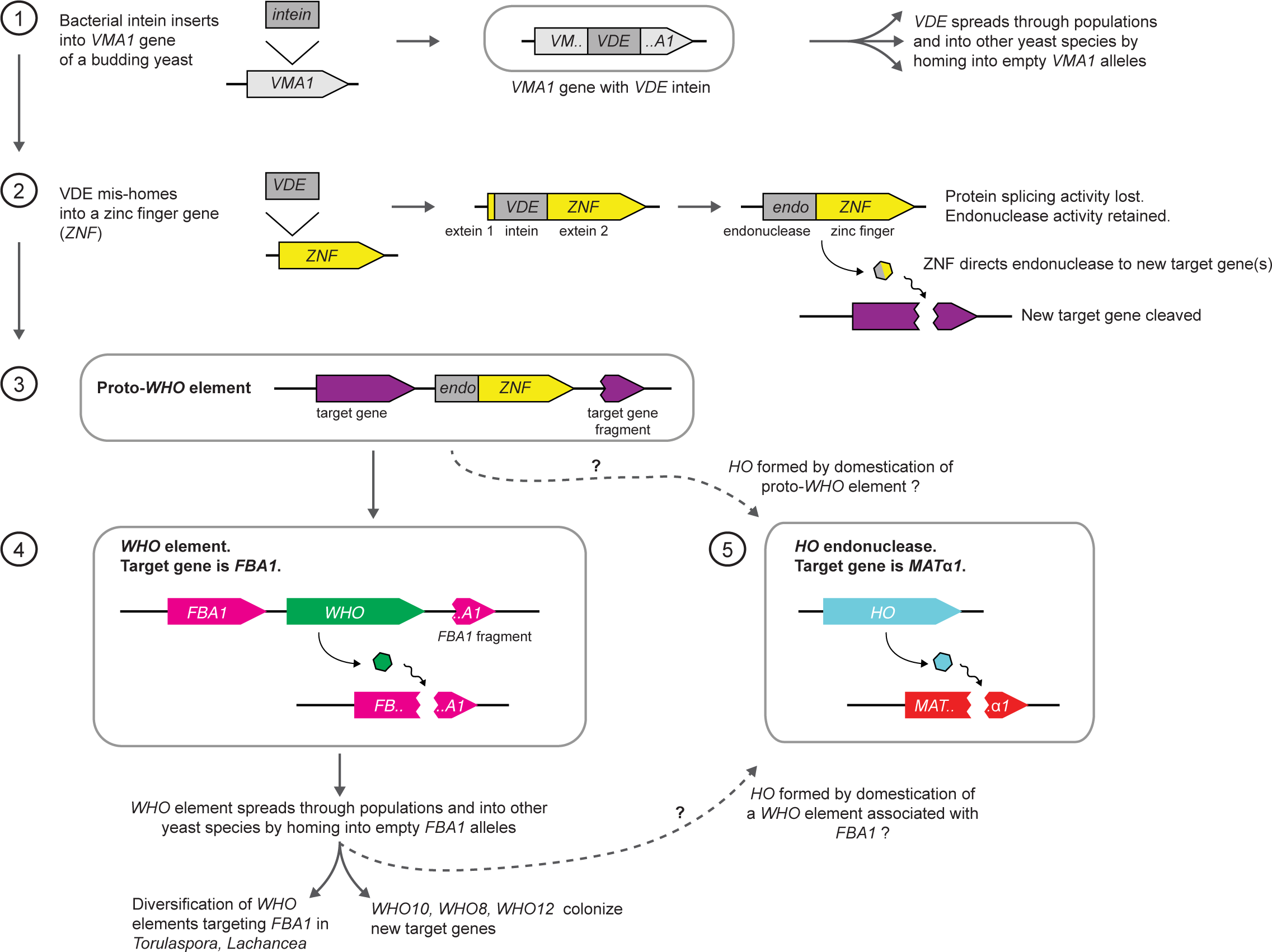
Model for the origin of *HO* by domestication of a *WHO* element. See Discussion for details. The dashed arrows indicate two possible routes to *HO*, from ancestral *WHO* elements that either were, or were not, specifically associated with *FBA1*.

The domestication of a *WHO* element to become *HO* is similar to the domestication of the transposon-derived genes *KAT1* and *α3* to act as generators of double-strand breaks at the *MAT* locus during mating-type switching in *K. lactis* (Barsoum et al., 2010; Rusche and Rine, 2010; Rajaei et al., 2014). In all three cases, a mobile element gene was domesticated in a genome that already had a three-locus arrangement and probably switched mating types by a passive process based on homologous recombination without a specific mechanism for making a double-strand break at *MAT*. Why were mobile genetic elements repeatedly recruited into these switching systems? If we consider that, in any population of haploid cells, (1) mating-type switching can only increase a cell’s probability of mating, (2) mating leads to the formation of a diploid and inevitably to meiosis, albeit many vegetative generations later, and (3) homing genetic elements can only home during meiosis, it logically follows that it is in a homing genetic element’s self-interest to increase the frequency of mating-type switching (Hanson and Wolfe, 2017). Thus, a *WHO* element could increase its rate of spread into empty *FBA1* alleles in a population, if its WHO protein developed a secondary activity of cleaving the *MAT* locus as well as cleaving *FBA1*. Since frequent and accurate switching are probably favored by natural selection (Hanson et al., 2014), the subsequent steps that domesticated a *WHO* element to form the non-mobile and exquisitely regulated gene *HO* (Stillman, 2013) would also have been advantageous.

## Acknowledgments

We thank Caroline Wilde, Weilong Hao, Warren Albertin and Robert Mans for strains, Devin Scannell and Mike Eisen for preliminary data on *T. globosa*, and Raúl Ortiz-Merino and Amanda Lohan for assistance. We thank Sara Hanson, Geraldine Butler and Wolfe lab members for comments on the manuscript. We thank Jo Dicks for permission to use the NCYC696 sequence data, which were produced by the UK National Collection of Yeast Cultures, in partnership with The Earlham Institute, using funding awarded to the Institute of Food Research, Norwich, UK by the Biotechnology and Biological Sciences Research Council. This study was supported by Science Foundation Ireland (13/IA/1910) and the European Research Council (789341).

## Author Contributions

A.Y.C. and K.H.W. designed the research; A.Y.C. constructed strains and plasmids; L.L. performed gene conversion assays; A.Y.C., S.B.-G., A.A.R.M., V.G., F.B. and S.D. sequenced genomes; K.P.B. performed bioinformatics analyses; A.Y.C. and K.H.W. wrote the paper.

## Materials and Methods

### Yeast isolates and genome sequences

New genome sequences for this study were obtained as follows. *T. delbrueckii* strains L09– L20 were from the strain collection of Lallemand Inc. (Montréal, Canada), generously provided by Dr. Caroline Wilde. They were sequenced at the University of Missouri core facility (Illumina, SE 1 x 50 bp). *T. delbrueckii* strain NCYC696 data was downloaded from <u>opendata.ifr.ac.uk/NCYC</u> on 23-Feb-2017 as unassembled Illumina sequence reads (PE 2 x 100 bp). *T. globosa* strains CBS764^T^ and CBS2947 were purchased from the Westerdijk Institute (Netherlands) and sequenced using both Illumina (PE 2 x 150 bp; BGI Tech Solutions, Hong Kong) and Pacific Biosciences Sequel technologies (1 SMRT cell; Earlham Institute, Norwich, UK). *T. pretoriensis* strain CBS2187^T^ (Illumina, PE 2 x 100 bp + 6 kb library MP 2 x 100 bp) and *T. franciscae* strain CBS2926^T^ (Illumina, PE 2 x 100 bp) were sequenced and assembled at INRAE Montpellier. Eight other *T. pretoriensis* strains (CBS2785, CBS5080, CBS9333, CBS11100, CBS11121, CBS11123, CBS11124 from the Westerdijk Institute, and UWOPS 81-1046.2 from M.A. Lachance, University of Western Ontario) were sequenced at the Earlham Institute using their proprietary LITE protocol for the Illumina platform. *Zygotorulaspora mrakii* strain NRRL Y-6702^T^ was obtained from the USDA Agricultural Research Service (Peoria, IL, USA) and sequenced at the Earlham Institute using both Pacific Biosciences RSII (4 SMRT cells) and Illumina LITE methods. *Naumovozyma castellii* strains Y056, Y174, Y287 and Y668 were gifts from Prof. Jure Piškur (Lund University, Sweden) and were sequenced at the Earlham Institute using the Illumina LITE method.

Cultures were grown under standard rich-medium conditions. DNA for Illumina sequencing was harvested from stationary-phase cultures by homogenization with glass beads followed by phenol-chloroform extraction and ethanol precipitation. Purified DNA was concentrated with the Genomic DNA Clean & Concentrator-10 (Zymo Research, catalog D4010). DNA for PacBio sequencing was prepared as in Ortiz-Merino et al. (2017).

Illumina data was assembled using SPAdes version v3.11.1 (Bankevich et al., 2012). PacBio data was assembled using HGAP3 (Chin et al., 2013).

Other genome sequences used in this study were taken from the NCBI database. The previously published genome sequences for *Torulaspora* species are from Gordon et al. (2011), Gomez-Angulo et al. (2015), Tondini et al. (2018), Galeote et al. (2018) and Shen et al. (2018), and for *Lachancea* species are from Souciet et al. (2009), Vakirlis et al. (2016), and Freel et al. (2016).

### Construction of S. cerevisiae ade2::TdFBA1 strains

*S. cerevisiae* strains in which the coding region (ORF) of *T. delbrueckii FBA1* was integrated into the *S. cerevisiae ADE2* gene, in opposite orientation to *ADE2* so that it is not transcribed, were constructed using CRISPR-Cas9 as follows. The *ADE2*-targeting sgRNA *ADE2.*Y from DiCarlo et al. (2013) was synthesized as a gene fragment by Integrated DNA Technologies, and inserted into the sgRNA plasmid pMEL13 from Mans et al. (2015) by restriction digestion and ligation. This plasmid was then transformed into *S. cerevisiae* strain IMX585 expressing Cas9 (Mans et al., 2015), together with a repair template containing the *delbrueckii FBA1* ORF (*TdFBA1-S* or *TdFBA1-R* allele) flanked with homology to *S. cerevisiae ADE2* in reverse orientation (bases 564456..564832 and 565952..566366 of *S. cerevisiae* chromosome XV). Transformants were selected on YPAD (YPD (Formedium) supplemented with 40 μg/ml adenine sulfate (Sigma)) containing 200 μg/ml G418. *ADE2* knockouts were identified by formation of red colonies. Successful integrants were confirmed by PCR amplification of the *ade2::TdFBA1* locus and Sanger sequencing (Eurofins). Two replicate *ade2::TdFBA1-S* strains were designated C1 and C4, and two replicate *ade2::TdFBA1-R* strains were designated L1 and L3. Sequences of the *ade2::TdFBA1-S* and *ade2::TdFBA1-R* constructs are given in Supplementary File S1.

### *WHO6* expression plasmid construction

Replicating plasmids constitutively expressing Who6, Who6-HA, or GFP-HA were constructed in the panARS replicating vector pIL75 (Liachko and Dunham, 2013) containing a *KanMX* marker. The nucleotide sequences of these plasmids (pWHO6, pWHO6-HA, pGFP-HA) are given in Supplementary File S1. In pWHO6, the *WHO6* gene from *T. delbrueckii* strain L09 was placed under the control of the promoter and terminator of the *T. delbrueckii* glyceraldehyde-3-phosphate dehydrogenase gene *TDH3* (*TDEL0E04750*). These regions were amplified by PCR from *T. delbrueckii* genomic DNA using high fidelity polymerase (New England Biolabs, M0492S) and inserted into pIL75 by restriction digestion and ligation. Plasmid pWHO6-HA is identical to pWHO6 except that its *WHO6* gene is fused to a C-terminal 3xHA tag. A similar control plasmid (pGFP-HA) was made containing a GFP gene fused to 3xHA tag (GBlock made by Integrated DNA Technologies and inserted into pIL75 by restriction digestion and ligation), under the control of the *T. delbrueckii TDH3* promoter and terminator.

### Gene conversion assays

The *S. cerevisiae ade2::TdFBA1* strains (C1, C4, L1, L3) as described above were transformed with the plasmids pWHO6, pWHO6-HA, or pGFP-HA. Multiple independent transformations were made to ensure that all gene conversion events recovered were independent. First, pWHO6-HA and pGFP-HA were each transformed into strains C1, C4, L1 and L3, and the whole genomes of these 8 strains were sequenced, resulting in the unexpected discovery of gene conversion at the *ade2::TdFBA1* locus when C1 and C4 were transformed with pWHO6-HA (called survivors 11 and 10, respectively, in Fig. 2B). Genome sequencing was done by BGI Tech Solutions using a BGISEQ instrument with 50-bp single-end reads. Second, pWHO6 (no 3xHA tag) and pGFP-HA were each transformed 10 times into strains C4 and L1. The *ade2::TdFBA1* locus from each transformant was amplified by PCR with a high-fidelity polymerase (New England Biolabs, M0492S) and Sanger sequenced. Primers for amplification were TGACCACGTTAATGGCTCC and CACCAGCTCCAGCGATAATTG.

For transformation, *S. cerevisiae* cells were grown overnight in liquid cultures of YPAD. Cultures were reinoculated in 50 ml and grown to mid-log phase. Cells were incubated with 1M LiAc, 50% PEG, salmon sperm DNA and plasmid DNA for 30 min, and then heat shocked at 42°C for 15 min. Transformants were selected on YPAD containing 200 μg/ml G418. Only one colony was picked from each transformation plate. The number of colonies obtained on the plates with the combination C4 + pWHO6 was dramatically lower than on the 3 other combinations. Individual colonies were grown in liquid YPAD overnight, before genomic DNA was harvested using a QIAamp DNA Mini kit (Qiagen).

## Supplemental Information

**Figure S1.**
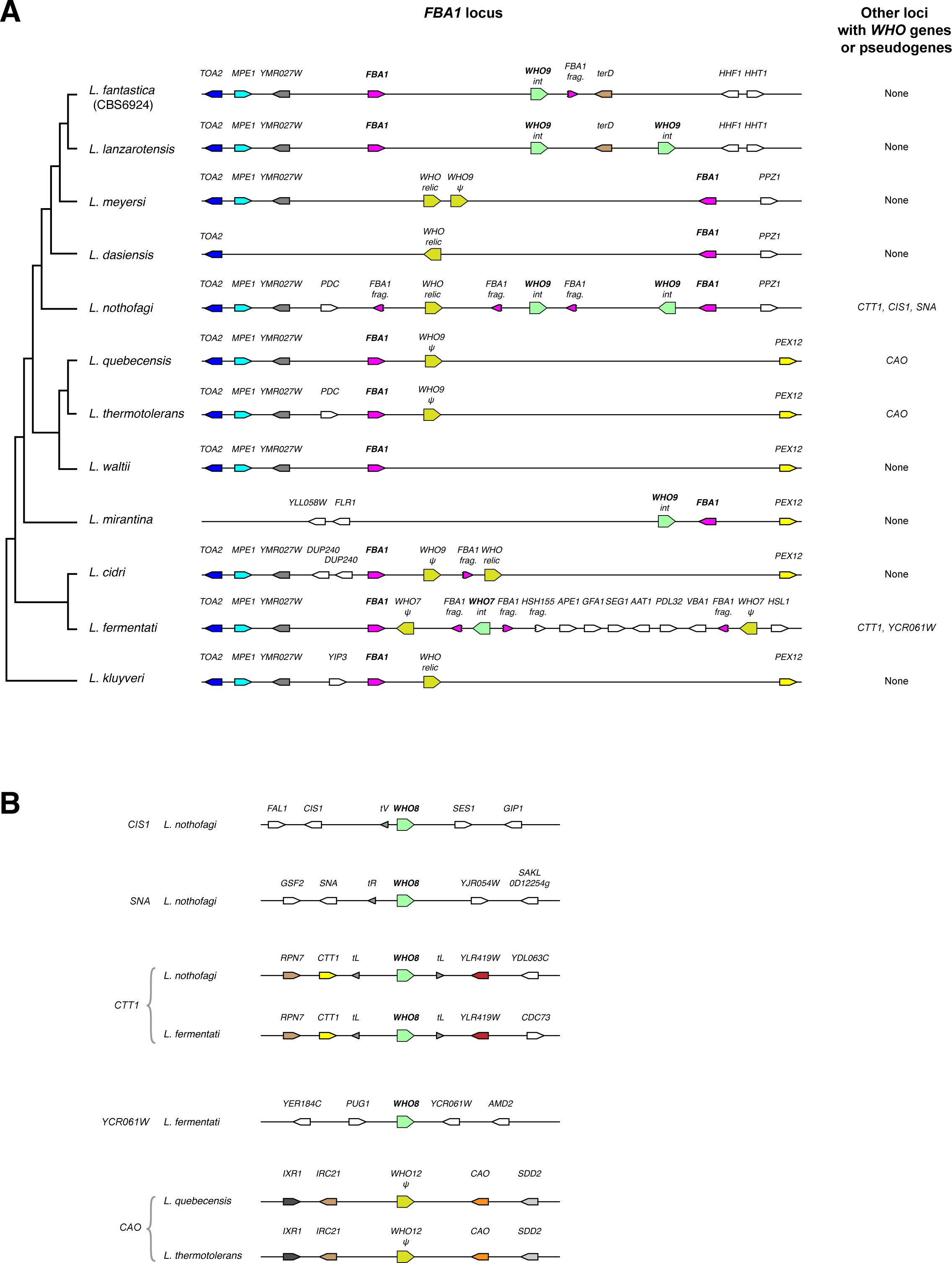
Genomic organization and domain structure of *WHO* genes in *Lachancea* species. **(A)** *WHO* genes and pseudogenes downstream of *FBA1* in *Lachancea* species. Fragments of the 3’ end of *FBA1* are marked. For *WHO* genes, “int” indicates an intact gene, “ψ” indicates a substantial pseudogene, and “relic” indicates a highly degraded pseudogene. Genomic views are schematic and not drawn to scale. The phylogenetic tree is based on Vakirlis et al. (2016). **(B)** All *WHO* genes at locations other than *FBA1* in *Lachancea* species.

**Figure S2.**
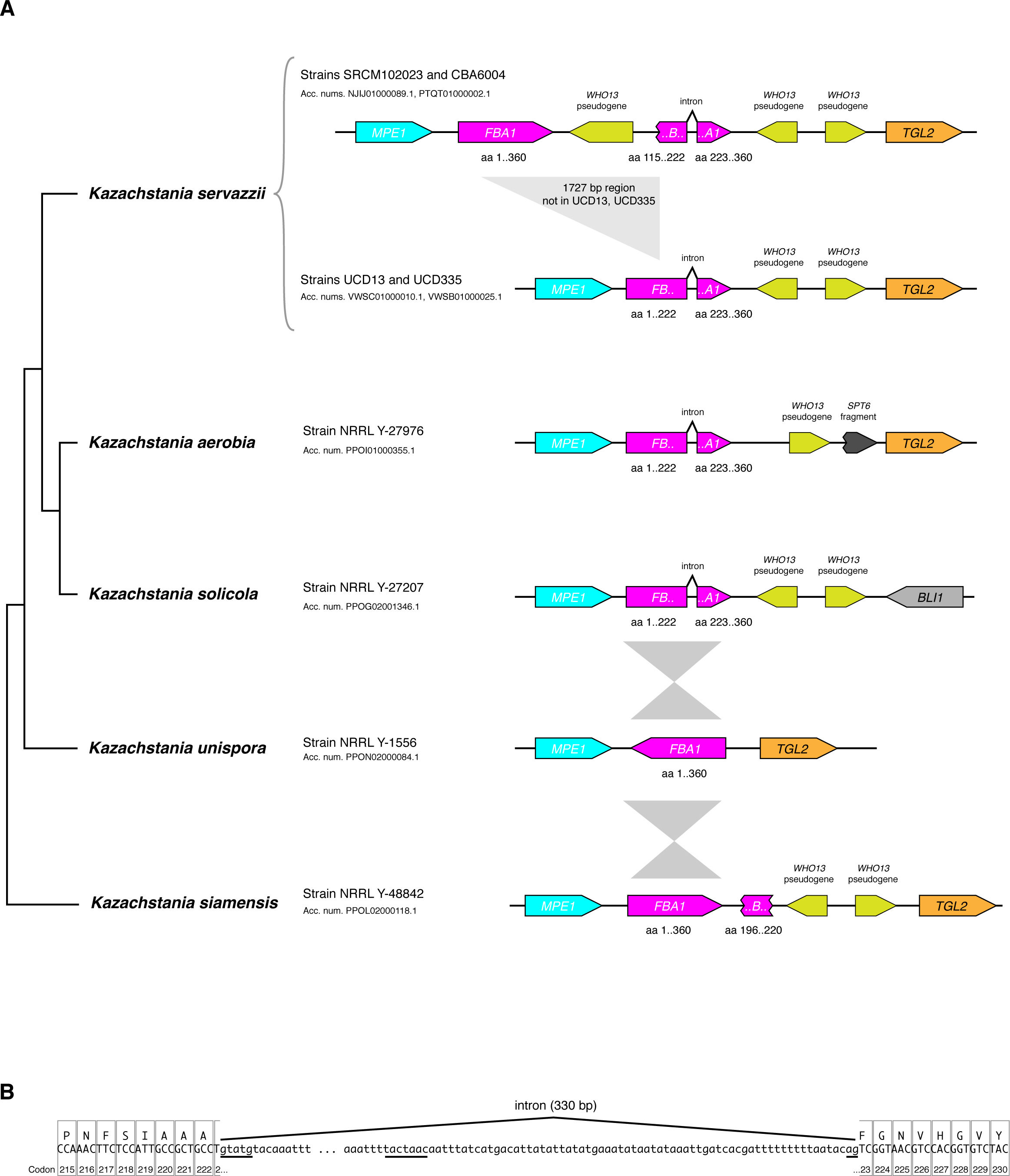
Some *Kazachstania* species have an intron in *FBA1* at a location corresponding to the *WHO* cleavage site in *Torulaspora*. **(A)** Genome organization around *FBA1* in *Kazachstania servazzii* and its close relatives. Data from 4 strains of *K. servazzii* is shown. *FBA1* genes, exons and fragments are colored magenta and their amino acid coordinates in the Fba1 protein are indicated. The phylogenetic tree topology is from Shen et al. (2018) and Vaughan-Martini et al. (2011). The genomes of 13 other species of *Kazachstania* that are outgroups to the ones shown here have no *FBA1* intron and no *WHO* genes (data from Shen et al., 2018). **(B)** Sequence of the intron in the *FBA1* gene of *K. servazzii* strain UCD13. The intron is located between the first and second bases of codon 223. Intron sequence is shown in lowercase, with exons and their translations in uppercase. Core intron sequence motifs are underlined.

**Supplementary File 1.** Sequences of the plasmids pWHO6, pWHO6-HA and pGFP-HA, and of the *ade2::TdFBA1-S* and *ade2::TdFBA1-R* constructs.

**Table S1.** T. delbrueckii genome sequence data used in this study.

| Strain name | *FBA1* region organization | *FBA1* gene sequence | NCBI accession number | Reference |
| --- | --- | --- | --- | --- |
| CBS1146<br>(type strain) | Shown in Fig. 1 | CBS1146 in Fig. 3 | HE616742-HE616749 | Gordon et al. (2011) |
| L09 | Shown in Fig. 1 | L09 in Fig. 3 | XXXXXX | This study |
| COFT1 | Same as L09 | Identical to L09 | CP027647-CP027655 | Tondini et al. (2018) |
| SRCM101298 | Same as L09 | Identical to L09 | NJII00000000 | S.H. Cho et al., published only in database |
| NRRL Y-50541 | Shown in Fig. 1 | NRRL Y-50541 in Fig. 3 | CP011778-CP011785 | Gomez-Angulo et al. (2015) |
| L16 | Shown in Fig. 1 | L16 in Fig. 3 | XXXXXX | This study |
| NCYC696 | Shown in Fig. 1 | NCYC696 in Fig. 3 | XXXXXX | NCYC / Earlham Institute |
| L18 | Shown in Fig. 1 | L18 in Fig. 3 | XXXXXX | This study |
| L11 | Same as L18 | L11 in Fig. 3 | XXXXXX | This study |
| L15 | Same as L18 | Identical to L11 | XXXXXX | This study |
| L19 | Same as L18 | Identical to L11 | XXXXXX | This study |
| L12 | Same as L18 | L12 in Fig. 3 | XXXXXX | This study |
| L13 | Same as L18 | Identical to L12 | XXXXXX | This study |
| L10 | Same as L18 | L10 in Fig. 3 | XXXXXX | This study |
| L20 | Same as L18 | L20 in Fig. 3 | XXXXXX | This study |

## References

Anraku, Y., Mizutani, R., and Satow, Y. (2005). Protein splicing: its discovery and structural insight into novel chemical mechanisms. IUBMB Life 57, 563–574.

Bakhrat, A., Baranes, K., Krichevsky, O., Rom, I., Schlenstedt, G., Pietrokovski, S., and Raveh, D. (2006). Nuclear import of ho endonuclease utilizes two nuclear localization signals and four importins of the ribosomal import system. J Biol Chem 281, 12218–12226.

Bakhrat, A., Jurica, M.S., Stoddard, B.L., and Raveh, D. (2004). Homology modeling and mutational analysis of Ho endonuclease of yeast. Genetics 166, 721–728.

Bankevich, A., Nurk, S., Antipov, D., Gurevich, A.A., Dvorkin, M., Kulikov, A.S., Lesin, V.M., Nikolenko, S.I., Pham, S., Prjibelski, A.D., et al. (2012). SPAdes: a new genome assembly algorithm and its applications to single-cell sequencing. J Comput Biol 19, 455–477.

Barsoum, E., Martinez, P., and Astrom, S.U. (2010). Alpha3, a transposable element that promotes host sexual reproduction. Genes Dev 24, 33–44.

Belfort, M. (2017). Mobile self-splicing introns and inteins as environmental sensors. Curr Opin Microbiol 38, 51–58.

Belfort, M., Stoddard, B.L., Wood, D.W., and Derbyshire, V., eds. (2005). Homing Endonucleases and Inteins (Berlin: Springer-Verlag).

Boles, E., and Zimmermann, F.K. (1993). *Saccharomyces cerevisiae* phosphoglucose isomerase and fructose bisphosphate aldolase can be replaced functionally by the corresponding enzymes of *Escherichia coli* and *Drosophila melanogaster*. Curr Genet 23, 187–191.

Burt, A., and Koufopanou, V. (2004). Homing endonuclease genes: the rise and fall and rise again of a selfish element. Curr Opin Genet Devel 14, 609–615.

Burt, A., and Trivers, R. (2008). Genes in conflict: The biology of selfish genetic elements (Harvard University Press).

Butler, G., Kenny, C., Fagan, A., Kurischko, C., Gaillardin, C., and Wolfe, K.H. (2004). Evolution of the *MAT* locus and its Ho endonuclease in yeast species. Proc Natl Acad Sci USA 101, 1632–1637.

Chevalier, B.S., and Stoddard, B.L. (2001). Homing endonucleases: structural and functional insight into the catalysts of intron/intein mobility. Nucleic Acids Res 29, 3757–3774.

Chin, C.S., Alexander, D.H., Marks, P., Klammer, A.A., Drake, J., Heiner, C., Clum, A., Copeland, A., Huddleston, J., Eichler, E.E., et al. (2013). Nonhybrid, finished microbial genome assemblies from long-read SMRT sequencing data. Nature Methods 10, 563–569.

Chiruvella, K.K., Rajaei, N., Jonna, V.R., Hofer, A., and Astrom, S.U. (2016). Biochemical Characterization of Kat1: a Domesticated hAT-Transposase that Induces DNA Hairpin Formation and MAT-Switching. Scientific reports 6, 21671.

Dalgaard, J.Z., Klar, A.J., Moser, M.J., Holley, W.R., Chatterjee, A., and Mian, I.S. (1997). Statistical modeling and analysis of the LAGLIDADG family of site-specific endonucleases and identification of an intein that encodes a site-specific endonuclease of the HNH family. Nucleic Acids Res 25, 4626–4638.

Dicarlo, J.E., Norville, J.E., Mali, P., Rios, X., Aach, J., and Church, G.M. (2013). Genome engineering in *Saccharomyces cerevisiae* using CRISPR-Cas systems. Nucleic Acids Res 41, 4336–4343.

Dickinson, J.R., and Williams, A.S. (1986). A genetic and biochemical analysis of the role of gluconeogenesis in sporulation of *Saccharomyces cerevisiae*. J Gen Microbiol 132, 2605–2610.

Fabre, E., Muller, H., Therizols, P., Lafontaine, I., Dujon, B., and Fairhead, C. (2005). Comparative genomics in hemiascomycete yeasts: evolution of sex, silencing and subtelomeres. Mol Biol Evol 22, 856–873.

Freel, K.C., Friedrich, A., Sarilar, V., Devillers, H., Neuveglise, C., and Schacherer, J. (2016). Whole-Genome Sequencing and Intraspecific Analysis of the Yeast Species *Lachancea quebecensis*. Genome Biol Evol 8, 733–741.

Galeote, V., Bigey, F., Devillers, H., Ortiz-Merino, R.A., Dequin, S., Wolfe, K.H., and Neuveglise, C. (2018). Genome Sequence of *Torulaspora microellipsoides* CLIB 830(T). Genome Announc 6, e00615–00618.

Gimble, F.S. (2000). Invasion of a multitude of genetic niches by mobile endonuclease genes. FEMS Microbiol Lett 185, 99–107.

Gimble, F.S., and Thorner, J. (1992). Homing of a DNA endonuclease gene by meiotic gene conversion in *Saccharomyces cerevisiae*. Nature 357, 301–306.

Gimble, F.S., and Thorner, J. (1993). Purification and characterization of VDE, a site-specific endonuclease from the yeast *Saccharomyces cerevisiae*. J Biol Chem 268, 21844–21853.

Gomez-Angulo, J., Vega-Alvarado, L., Escalante-Garcia, Z., Grande, R., Gschaedler-Mathis, A., Amaya-Delgado, L., Arrizon, J., and Sanchez-Flores, A. (2015). Genome Sequence of *Torulaspora delbrueckii* NRRL Y-50541, Isolated from Mezcal Fermentation. Genome Announc 3.

Gordon, J.L., Armisen, D., Proux-Wera, E., OhEigeartaigh, S.S., Byrne, K.P., and Wolfe, K.H. (2011). Evolutionary erosion of yeast sex chromosomes by mating-type switching accidents. Proc Natl Acad Sci USA 108, 20024–20029.

Gouy, M., Guindon, S., and Gascuel, O. (2010). SeaView version 4: A multiplatform graphical user interface for sequence alignment and phylogenetic tree building. Mol Biol Evol 27, 221–224.

Green, C.M., Novikova, O., and Belfort, M. (2018). The dynamic intein landscape of eukaryotes. Mobile DNA 9, 4.

Haber, J.E. (2016). A Life Investigating Pathways That Repair Broken Chromosomes. Annu Rev Genet 50, 1–28.

Haber, J.E., and Wolfe, K. (2005). Evolution and function of HO and VDE endonucleases in fungi. In Homing Endonucleases and Inteins, M. Belfort, B. Stoddard, D. Wood, and V. Derbyshire, eds. (Berlin: Springer-Verlag), pp. 161–176.

Hanson, S.J., Byrne, K.P., and Wolfe, K.H. (2014). Mating-type switching by chromosomal inversion in methylotrophic yeasts suggests an origin for the three-locus *Saccharomyces cerevisiae* system. Proc Natl Acad Sci USA 111, E4851–4858.

Hanson, S.J., and Wolfe, K.H. (2017). An evolutionary perspective on yeast mating-type switching. Genetics 206, 9–32.

Herskowitz, I. (1988). Life cycle of the budding yeast *Saccharomyces cerevisiae*. Microbiol Rev 52, 536–553.

Huang, S., Tao, X., Yuan, S., Zhang, Y., Li, P., Beilinson, H.A., Zhang, Y., Yu, W., Pontarotti, P., Escriva, H., et al. (2016). Discovery of an Active RAG Transposon Illuminates the Origins of V(D)J Recombination. Cell 166, 102–114.

Jin, Y., Binkowski, G., Simon, L.D., and Norris, D. (1997). Ho endonuclease cleaves *MAT* DNA in vitro by an inefficient stoichiometric reaction mechanism. J Biol Chem 272, 7352–7359.

Kaiser, B.K., Clifton, M.C., Shen, B.W., and Stoddard, B.L. (2009). The structure of a bacterial DUF199/WhiA protein: domestication of an invasive endonuclease. Structure 17, 1368–1376.

Keeling, P.J., and Roger, A.J. (1995). The selfish pursuit of sex. Nature 375, 283.

Kostriken, R., Strathern, J.N., Klar, A.J., Hicks, J.B., and Heffron, F. (1983). A site-specific endonuclease essential for mating-type switching in *Saccharomyces cerevisiae*. Cell 35, 167–174.

Koufopanou, V., and Burt, A. (2005). Degeneration and domestication of a selfish gene in yeast: molecular evolution versus site-directed mutagenesis. Mol Biol Evol 22, 1535–1538.

Koufopanou, V., Goddard, M.R., and Burt, A. (2002). Adaptation for horizontal transfer in a homing endonuclease. Mol Biol Evol 19, 239–246.

Krassowski, T., Kominek, J., Shen, X.X., Opulente, D.A., Zhou, X., Rokas, A., Hittinger, C.T., and Wolfe, K.H. (2019). Multiple Reinventions of Mating-type Switching during Budding Yeast Evolution. Curr Biol 29, 2555–2562.

Kurtzman, C.P. (2011). *Torulaspora* Lindner (1904). In The Yeasts, A Taxonomic Study, C.P. Kurtzman, J.W. Fell, and T. Boekhout, eds. (Amsterdam: Elsevier), pp. 867–874.

Lee, C.S., and Haber, J.E. (2015). Mating-type gene switching in *Saccharomyces cerevisiae*. Microbiology Spectrum 3, MDNA3-0013-2014.

Li, W., Tucker, A.E., Sung, W., Thomas, W.K., and Lynch, M. (2009). Extensive, recent intron gains in *Daphnia* populations. Science 326, 1260–1262.

Liachko, I., and Dunham, M.J. (2013). An Autonomously Replicating Sequence for use in a wide range of budding yeasts. FEMS Yeast Res 14, 364–367.

Lobo, Z. (1984). *Saccharomyces cerevisiae* aldolase mutants. J Bacteriol 160, 222–226.

Mans, R., van Rossum, H.M., Wijsman, M., Backx, A., Kuijpers, N.G., van den Broek, M., Daran-Lapujade, P., Pronk, J.T., van Maris, A.J., and Daran, J.M. (2015). CRISPR/Cas9: a molecular Swiss army knife for simultaneous introduction of multiple genetic modifications in *Saccharomyces cerevisiae*. FEMS Yeast Res 15.

McDowell, J.M., and Meyers, B.C. (2013). A transposable element is domesticated for service in the plant immune system. Proc Natl Acad Sci USA 110, 14821–14822.

Meiron, H., Nahon, E., and Raveh, D. (1995). Identification of the heterothallic mutation in HO-endonuclease of *S. cerevisiae* using *HO/ho* chimeric genes. Curr Genet 28, 367–373.

Moore, J.K., and Haber, J.E. (1996). Cell cycle and genetic requirements of two pathways of nonhomologous end-joining repair of double-strand breaks in *Saccharomyces cerevisiae*. Mol Cell Biol 16, 2164–2173.

Motl, J.A., and Chalker, D.L. (2009). Subtraction by addition: domesticated transposases in programmed DNA elimination. Genes Dev 23, 2455–2460.

Moure, C.M., Gimble, F.S., and Quiocho, F.A. (2002). Crystal structure of the intein homing endonuclease PI-SceI bound to its recognition sequence. Nature Struct Biol 9, 764–770.

Muller, H., Hennequin, C., Dujon, B., and Fairhead, C. (2007). Ascomycetes: the *Candida MAT* locus: Comparing *MAT* in the genomes of hemiascomycetous yeasts In Sex in Fungi, J. Heitman, J.W. Kronstad, J.W. Taylor, and L.A. Casselton, eds. (Washington, D.C.: ASM Press), pp. 247–263.

Munchow, S., Ferring, D., Kahlina, K., and Jansen, R.P. (2002). Characterization of *Candida albicans ASH1* in *Saccharomyces cerevisiae*. Curr Genet 41, 73–81.

Neiman, A.M. (2005). Ascospore Formation in the yeast *Saccharomyces cerevisiae*. Microbiol Mol Biol Rev 69, 565–584.

Nickoloff, J.A., Singer, J.D., and Heffron, F. (1990). In vivo analysis of the *Saccharomyces cerevisiae* HO nuclease recognition site by site-directed mutagenesis. Mol Cell Biol 10, 1174–1179.

Novikova, O., Topilina, N., and Belfort, M. (2014). Enigmatic distribution, evolution, and function of inteins. J Biol Chem 289, 14490–14497.

Okuda, Y., Sasaki, D., Nogami, S., Kaneko, Y., Ohya, Y., and Anraku, Y. (2003). Occurrence, horizontal transfer and degeneration of VDE intein family in Saccharomycete yeasts. Yeast 20, 563–573.

Ortiz-Merino, R.A., Kuanyshev, N., Braun-Galleani, S., Byrne, K.P., Porro, D., Branduardi, P., and Wolfe, K.H. (2017). Evolutionary restoration of fertility in an interspecies hybrid yeast, by whole-genome duplication after a failed mating-type switch. PLoS Biol 15, e2002128.

Oshima, Y. (1993). Homothallism, mating-type switching, and the controlling element model in *Saccharomyces cerevisiae*. In The Early Days of Yeast Genetics, M.N. Hall, and P. Linder, eds. (New York: Cold Spring Harbor Laboratory Press), pp. 291–304.

Pietrokovski, S. (1994). Conserved sequence features of inteins (protein introns) and their use in identifying new inteins and related proteins. Protein Sci 3, 2340–2350.

Poulter, R.T., Goodwin, T.J., and Butler, M.I. (2007). The nuclear-encoded inteins of fungi. Fungal Genet Biol 44, 153–179.

Rajaei, N., Chiruvella, K.K., Lin, F., and Astrom, S.U. (2014). Domesticated transposase Kat1 and its fossil imprints induce sexual differentiation in yeast. Proc Natl Acad Sci USA 111, 15491–15496.

Rusche, L.N., and Rine, J. (2010). Switching the mechanism of mating type switching: a domesticated transposase supplants a domesticated homing endonuclease. Genes Dev 24, 10–14.

Russell, D.W., Jensen, R., Zoller, M.J., Burke, J., Errede, B., Smith, M., and Herskowitz, I. (1986). Structure of the *Saccharomyces cerevisiae HO* gene and analysis of its upstream regulatory region. Mol Cell Biol 6, 4281–4294.

Scazzocchio, C. (2000). The fungal GATA factors. Curr Opin Microbiol 3, 126–131.

Schwelberger, H.G., Kohlwein, S.D., and Paltauf, F. (1989). Molecular cloning, primary structure and disruption of the structural gene of aldolase from *Saccharomyces cerevisiae*. Eur J Biochem 180, 301–308.

Shen, X.X., Opulente, D.A., Kominek, J., Zhou, X., Steenwyk, J.L., Buh, K.V., Haase, M.A.B., Wisecaver, J.H., Wang, M., Doering, D.T., et al. (2018). Tempo and mode of genome evolution in the budding yeast subphylum. Cell 175, 1533–1545 e1520.

Souciet, J.L., Dujon, B., Gaillardin, C., Johnston, M., Baret, P.V., Cliften, P., Sherman, D.J., Weissenbach, J., Westhof, E., Wincker, P., et al. (2009). Comparative genomics of protoploid Saccharomycetaceae. Genome Res 19, 1696–1709.

Stillman, D.J. (2013). Dancing the cell cycle two-step: regulation of yeast G1-cell-cycle genes by chromatin structure. Trends Biochem Sci 38, 467–475.

Taylor, G.K., Petrucci, L.H., Lambert, A.R., Baxter, S.K., Jarjour, J., and Stoddard, B.L. (2012). LAHEDES: the LAGLIDADG homing endonuclease database and engineering server. Nucleic Acids Res 40, W110–116.

Tondini, F., Jiranek, V., Grbin, P.R., and Onetto, C.A. (2018). Genome Sequence of Australian Indigenous Wine Yeast *Torulaspora delbrueckii* COFT1 Using Nanopore Sequencing. Genome Announc 6.

Trifinopoulos, J., Nguyen, L.T., von Haeseler, A., and Minh, B.Q. (2016). W-IQ-TREE: a fast online phylogenetic tool for maximum likelihood analysis. Nucleic Acids Res 44, W232–235.

Vakirlis, N., Sarilar, V., Drillon, G., Fleiss, A., Agier, N., Meyniel, J.P., Blanpain, L., Carbone, A., Devillers, H., Dubois, K., et al. (2016). Reconstruction of ancestral chromosome architecture and gene repertoire reveals principles of genome evolution in a model yeast genus. Genome Res 26, 918–932.

Vaughan-Martini, A., Lachance, M.A., and Kurtzman, C.P. (2011). Kazachstania Zubkova (1971). In The Yeasts, a Taxonomic Study, C.P. Kurtzman, J.W. Fell, and T. Boekhout, eds. (Amsterdam: Elsevier), pp. 439–470.

Volff, J.N. (2006). Turning junk into gold: domestication of transposable elements and the creation of new genes in eukaryotes. Bioessays 28, 913–922.

Winge, O., and Roberts, C. (1949). A gene for diploidization in yeasts. C R Trav Lab Carlsberg Ser Physiol 24, 341–346.

